# Characterization of artificial riboswitches for Coxsackievirus B3 detection using machine learning

**DOI:** 10.64898/2026.10.04.756577

**Authors:** Jonathan Auyong, Hainian Audrey Long, Sonya Hu, Joel Jacob, Kristy Chan, Bhoomika Gupta, Atnasia Mengistu, Miyuki Tokuhara, Leila Khatib, William B. Andreopoulos

**Author notes:** These authors contributed equally to this work (Jonathan Auyong, Hainian Audrey Long, Sonya Hu, Joel Jacob, Kristy Chan, Bhoomika Gupta, Atnasia Mengistu, Miyuki Tokuhara).

## Abstract

With the increase of recycled water to offset water demand, the potential possibility to spread contagious RNA viruses, such as Coxsackievirus B3, increases. However, detection of viral particles remains challenging because of low viral concentrations in wastewater and high mutation rates of the RNA virus. Robust monitoring is needed with low cost and low infrastructure technologies to increase accessibility of monitoring technologies worldwide. Novel viral detection methods have been developed, based on synthetic riboswitches that bind to the target virus and trigger a reporter gene, thus amplifying the detection signals. Such monitoring technologies can leverage machine learning to optimize the candidate nucleic acid sequences used for detection. To support the design of effective riboswitches, we present a machine learning model for classifying riboswitch performance, integrating RNA sequence data with secondary structural features based on free energy calculations and parameters that represent single strandedness. This model uses a sparsely gated Mixture of Experts (MoE) architecture to route sequence and thermodynamic features to specialized experts, achieving strong generalization performance across cross-validation and held-out testing. When evaluated against baseline Decision Tree, Random Forest, Gaussian Naive Bayes, and Dense Neural Network classifiers, the MoE model demonstrated near-zero classification errors. Furthermore, post-hoc feature importance and k-mer analyzes reveal that near-perfect predictive performance in silico is strongly driven by sequence constructs in addition to generalizable folding mechanics. An ablation study of numerical feature and sequence inputs showed that while the Dense Neural Network also achieved high accuracy across all ablation conditions, the MoE architecture was retained as the primary model because its specialized subnetworks do not require all model parameters to be activated offering a potential computational advantage.

## 1 Introduction and Motivation

Climate change, increasing populations, urbanization, as well as data centers, contribute to water demands that reduce sustainable systems. Therefore, recycled water serves as an alternative water source to supplement water portfolios. By 2030, the water demand to cool data centers per person will exceed the drinking water requirement per person in Europe [1, 2]. Incorporating recycled water into water planning may also increase the exposure of enteric viruses such as Coxsackievirus due their stability through a waste treatment system [3]. Despite water use efficiency efforts for data centers, reducing municipal water sources will be needed for sustainable supplies which includes planning for the use of recycled water [4]. Current cooling technologies for data centers, such as evaporative cooling and cooling towers, may also exacerbate the exposure of enteric viruses such as EV 68 or EV 70 through aerosolization of the recycled water. As data centers increase in numbers worldwide, more robust, low cost, and low labor approaches are needed to monitor the presence of enteric viruses. Current monitoring technologies either require hours or days to produce results or require a commitment to costly infrastructure or equipment. As not all countries may have access to the resources necessary for virus detection, alternative methods are needed. Riboswitches have been demonstrated to detect viruses in less than 1 hour and using low cost technologies [5–7]. Riboswitches are regulatory sequences present in bacteria. They are composed of a ligand binding region, secondary structure which contains a ribosomal binding site. When the ligand binds to the binding region of the riboswitch, the secondary structure destabilizes and the conformational change linearizes the secondary structure. The RNA polymerase can bind to the ribosomal binding site and express the corresponding gene. Under thermodynamically favorable conditions, the riboswitch remains in a secondary structure thus turning the expression of the gene “off.” When a ligand matches the activation energy of the ligand binding regions, the conformational change occurs and the gene is turned “on.” [8].

However, the development of riboswitches for detection of viruses was based on trial and error and were labor intensive due to screening a large number of riboswitches to determine the optimal sequence [9]. In this paper, we examine iterative changes to the ligand binding region of artificial riboswitches and the corresponding free energy calculation for each iterative change to computational characterize artificial riboswitches. To support the design of effective riboswitches, we present a machine learning model for classifying riboswitch performance, integrating RNA sequence data with secondary structural features. These measurements represent indicators for the optimal riboswitch sequences.

## 2 Dataset Description

### 2.1 Background

For this study, the individual components of the riboswitch complex were first identified, extracted, and then assembled to form a complete construct. The sequence of the engineered riboswitch complex is composed of a 30 bp target sequence, a 30 bp trigger sequence, a published riboswitch sequence of varied length with a 20 bp linker sequence included, and a 10 bp reporter gene M-cherry. The trigger of the riboswitch is the most important component of the complex as it scans through the entire target region and identifies the correct target binding location. The trigger sequence was designed by shifting a 30 bp frame across the CVB3 RNA polymerase loci in a stepwise / base-pair-by-base-pair manner. The trigger sequence is reverse complementary to the RNA polymerase loci of the target sequence. The complete target sequence of Human Coxsackievirus B3 mRNA was downloaded from NCBI and then extracted for the RNA polymerase loci at 1879-2030 bp [42]. Both the riboswitch sequence and the linker sequence were extracted from the de novo-designed toehold switches from the study by Green et al. [43]. These riboswitches, also referred to as toehold switches by the authors, can be forward engineered above 400 bp, making them suitable for use as riboregulators for the purpose of detecting CVB3 virus.

The reporter gene M-cherry (synthetic construct monomeric red fluorescent protein gene) of the riboswitch was downloaded from NCBI and then extracted to use the first 10 bp [44]. Figure 1 highlights the architecture of the artificial riboswitch construct used in this study. The construct consists of trigger sequence, ribosome binding site (RBS), linker, and reporter gene. Upon binding of the complementary target RNA, the RBS becomes accessible, which enables for downstream reporter gene expression. Table 1 defines the terminology used in the synthetic riboswitch analysis.

**Table 1:** Summary of each component of the synthetic riboswitch.

| Component | Description |
| --- | --- |
| Target | Single-stranded RNA viral sequence for detection. |
| Trigger | Synthesized sequence that is reverse complementary to the target, designed to bind to the target. |
| Riboswitch | Secondary structure that unfolds when the trigger binds to the target. |
| Linker | Sequence that connect riboswitch and reporter. |
| Reporter | Fluorescent protein when expressed signals the presence of the target. |

**Figure 1:**
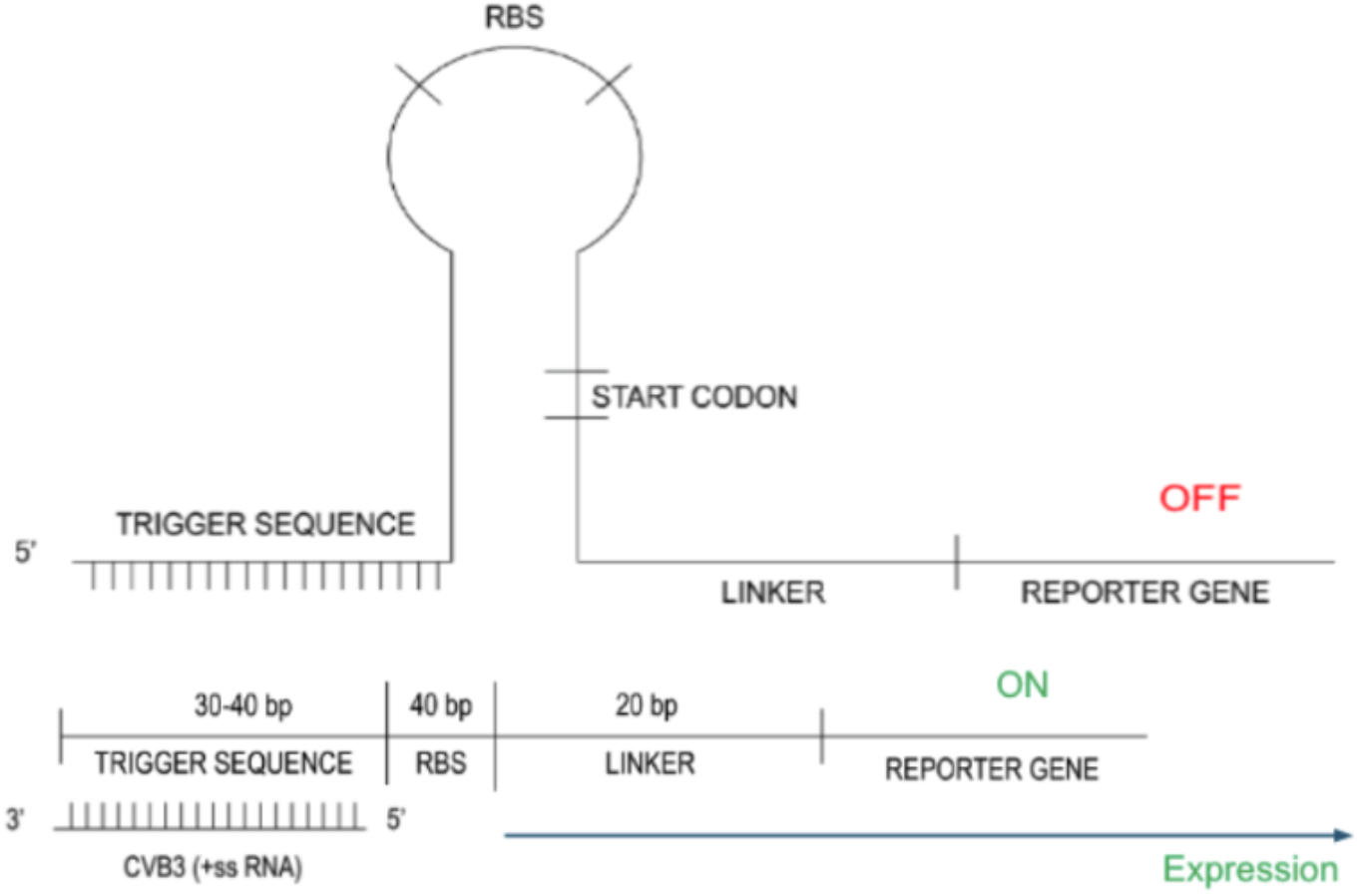
Architecture of artificial riboswitch construct in the OFF state (top) vs. ON state (bottom).

#### 2.1.1 Data Source

The engineered riboswitch constructs analyzed in this study consist of four primary sequence components: a target sequence, trigger sequence, riboswitch sequence, and a reporter gene (Figure 2). Each construct was assembled by combining a 30 bp target sequence, 30 bp trigger sequence, a published riboswitch sequence containing a 20 bp linker region, and a 10 bp fragment of the M-cherry reporter gene.

**Figure 2:**
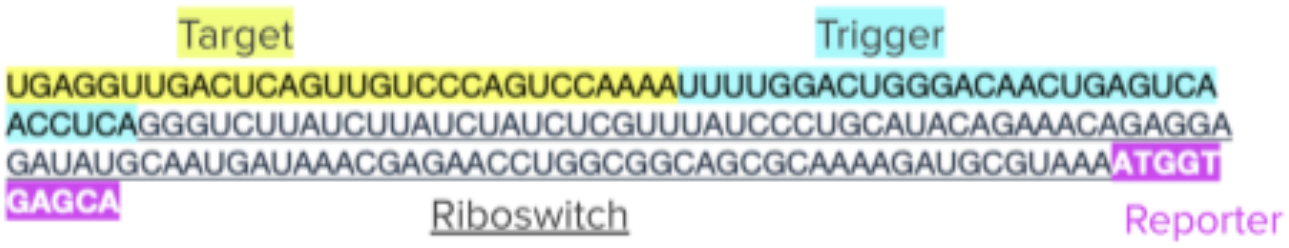
Example of a riboswitch complex sequence. Yellow: target. Blue: trigger. Underline: riboswitch. Bold: reporter.

The target sequence was derived from the RNA polymerase locus (positions 1879-2030 bp) of the Human Coxsackievirus B3 (CVB3) genome downloaded from the National Center for Biotechnology Information (NCBI) [42]. Trigger sequences were generated by sliding a 30 bp window across the target region one nucleotide at a time, producing reverse-complementary trigger sequences corresponding to each target location.

The riboswitch and linker sequences were obtained from the de novo-designed toehold switches reported by Green et al. [43], which function as programmable RNA regulatory elements that undergo conformational changes upon trigger binding to regulate downstream translation [17]. The reporter sequence was obtained from the synthetic monomeric red fluorescent protein (M-cherry) construct available through NCBI, from which the first 10 bp were extracted for incorporation into the riboswitch construct [44].

### 2.1.2 Data Generation

For each engineered riboswitch construct, the riboswitch, target, reporter, and complete complex sequences were analyzed using the ViennaRNA package [29] to predict RNA secondary structure and thermodynamic properties. ViennaRNA predicts RNA folding using thermodynamic models and partition function calculations [29, 31, 32]. Minimum free energy (MFE) values represent the thermodynamic stability of predicted RNA secondary structures, with more negative values indicating greater structural stability. Structural prediction generated multiple numerical features describing RNA folding and thermodynamic stability, including minimum free energy, GC and AT content, ensemble defect, single-strandedness, trigger-target binding percentage, loop energy, salt molarity, temperature, and structural properties for the riboswitch, target, reporter, and complete complex. These features provide quantitative descriptors of RNA stability and secondary structure that were subsequently used as inputs for machine learning models. Prediction results for each construct were exported as CSV files and then combined into a single dataset for downstream machine learning analysis.

### 2.2 Dataset Positive and Negative Class Labels

Each artificial riboswitch construct was assigned a binary label based on the pairing relationship between its trigger sequence and the corresponding target window within the Coxsackievirus B3 (CVB3) RNA-dependent RNA polymerase locus. Constructs were labeled positive when the trigger sequence was designed to be complementary to its assigned target window and was therefore expected to anneal correctly within this locus. Constructs were labeled negative when the trigger sequence was paired with a noncorresponding target window, producing an intended trigger-target mismatch. Thus, the negative class represents incorrect pairing within the CVB3 design space rather than a distinct non-CVB3 viral class.

This labeling scheme defines the machine-learning task as the classification of correct versus incorrect trigger-target pairings within the same viral target region. Both positive and negative constructs were derived from CVB3 target sequences and spanned the same set of 152 unique target-window locations. The labels were also reflected in the trigger-target binding percentage, which averaged 99.4% for positive constructs and 24.9% for negative constructs, consistent with stronger predicted annealing in correctly paired constructs and weaker predicted annealing in mismatched constructs.

The complete dataset contained 257,032 artificial riboswitch constructs represented by 54 sequence-derived, thermodynamic, structural, and experimental-condition columns generated through ViennaRNA secondary-structure prediction. Following label assignment, 27,512 constructs were labeled positive and 229,520 were labeled negative. To mitigate class imbalance during model training, 27,512 negative constructs were randomly sampled to match the number of positive constructs, producing a balanced dataset of 55,024 constructs with equal positive and negative class sizes.

A class-associated difference was also observed in the riboswitch design-generation metadata. The negative class was drawn from the “forward engineered 4” design generation, whereas the positive class included constructs from “144 orthogonal first gen”, “24 first gen”, and “forward engineered 4”. Because this metadata difference is not part of the intended trigger-target pairing definition, it was treated as a potential dataset confound in the interpretation of model performance.

### 2.3 Data Preprocessing

#### 2.3.1 Data Cleaning

Prior to machine learning analysis, features incompatible with the learning algorithms were removed from the dataset. Dot-bracket RNA secondary structure strings were removed because conventional machine learning algorithms cannot directly process symbolic sequence representations. Dictionary-based k-mer count outputs were excluded because they require additional feature engineering before numerical modeling. Features containing sample identifiers or metadata unrelated to riboswitch structure were removed to prevent noninformative variables from influencing model performance. Metadata fields unrelated to structural prediction, including sample identifiers and local information, were excluded to reduce noise and improve model performance.

#### 2.3.2 Feature Selection

Following preprocessing, the original 54-column dataset was reduced to a retained model-input set composed of 26 scalar numerical features and one sequence-valued feature, complex_nucleotides. The numerical features were derived from ViennaRNA structural and thermodynamic predictions, whereas complex_nucleotides represented the full riboswitch construct sequence. Accordingly, the apparent change from 27 retained inputs to 26 numerical inputs reflects a distinction between feature type rather than the removal of an additional feature: the 27th retained input was processed separately as sequence data. This sequence-valued feature was subsequently converted into an 840-dimensional one-hot encoded representation, corresponding to 210 nucleotide positions with four encoding dimensions per position. Thus, each construct is represented by 866 total model inputs: 26 numerical features and 840 sequence-derived features. This combined representation was provided to all five classification models: Decision Tree, Random Forest, Dense Neural Network, Gaussian Naive Bayes, and Mixture of Experts.

#### 2.3.3 Feature Encoding

In order to enable machine learning analysis of nucleotide sequences, one-hot encoding is applied to the riboswitch complex sequence [14]. Each nucleotide was represented as a four-dimensional binary vector, where adenine (A), cytosine (C), guanine (G), and uracil (U) were encoded as [1,0,0,0], [0,1,0,0], [0,0,1,0], and [0,0,0,1], respectively. Thymine (T) nucleotides originating from the reporter sequence were encoded identically to uracil (U), since both represent equivalent pyrimidine bases in this encoding scheme. Riboswitch complex sequences varied in length and were therefore standardized to a fixed length of 210 nucleotides by right-padding shorter sequences with the placeholder nucleotide “N”, represented by the null vector [0,0,0,0]. The padding strategy preserves the original sequence while providing a uniform input size required for downstream machine learning models [13]. Following one-hot encoding, each riboswitch complex sequence was represented as an 840-dimensional feature vector (210 × 4). Combined with the 26 numerical ViennaRNA-derived features, with each riboswitch construct represented by a total of 866 input features for model training.

#### 2.3.4 Feature Scaling

Numerical features were standardized using StandardScaler implementation from scikit-learn library [46]. Standardization transformed each feature to have zero mean and unit variance, preventing features with larger numerical ranges from dominating model training. The scaler was fitted using only the training data within each cross-validation fold and subsequently applied to the corresponding validation and test sets to prevent information leakage.

### 2.4 Limitations of data

A central limitation of the current dataset is that positive and negative labels are generated from a computational trigger-target pairing rule rather than from experimentally measured riboswitch activation. For a given trigger, only the correctly matched target window is labeled positive; mismatched target windows are labeled negative. This construction creates many near-neighbor sequences and can make the classification problem easier than a prospective biological design task. Future datasets should include experimentally validated riboswitch activity measurements and additional viral targets, such as poliovirus, MS2, enterovirus, and other RNA viruses. Cross-target and cross-virus evaluation would make it possible to distinguish genuine biological generalization from performance driven by repeated sequence windows, design-generation imbalance, or dataset-specific artifacts.

### 2.5 Machine learning methods for Classification

Five machine learning models were evaluated for predicting riboswitch complex stability: a Decision Tree, Random Forest, a Dense Neural Network (multilayer perceptron (MLP)), Gaussian Naive Bayes, and a Mixture of Experts (MoE) architecture. All models were trained using the same processed feature matrix and compared using identical training, validation, and evaluation procedures. For all five models, the input consisted of the 26 numerical features together with the 840-dimensional one-hot encoded sequence representation unless otherwise specified in the ablation experiments.

#### 2.5.1 Decision Tree

A Decision Tree classifier was implemented using scikit-learn’s DecisionTreeClassifier as a classical machine learning baseline [65]. Decision trees partition the feature space recursively by selecting decision rules to minimize prediction errors at each node. At each split, the algorithm evaluates all possible features and thresholds, choosing the one that best reduces the variance in the target variable. This process continues until a stopping criterion is met (e.g., maximum tree depth or minimum samples per leaf). Decision trees are interpretable, as the path from root to leaf represents a human-readable set of conditional rules. However, single decision trees are prone to overfitting and usually underperform on complex datasets. The model was configured with no maximum depth constraint, `min_samples_split = 2`, `min_samples_leaf = 1`, and `max_features = ‘sqrt’` to select a random subset of 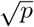 features at each split. The same preprocessing pipeline, train-test split, and 5-fold cross-validation protocol were applied as described above. In addition to classification performance, feature importances were extracted from the trained decision tree to assess which features contributed most to the classification decisions, providing an interpretability advantage over the neural network models.

#### 2.5.2 Random Forest

A Random Forest classifier was implemented using scikit-learn’s RandomForestClassifier as an ensemble-based baseline. The model consisted of 300 decision trees with max_features = ‘sqrt’, allowing each tree to consider a random subset of input features at each split. The predictions were obtained by aggregating the decisions across the ensemble [71].

#### 2.5.3 Dense Neural Network

A Dense Neural Network classifier was implemented as a deep learning baseline to evaluate classification performance without expert routing. The architecture consists of fully connected feedforward layers with LayerNorm, LeakyReLU activations, and Dropout (*p* = 0.3) to prevent overfitting on the 866-dimensional input feature space. The network was trained for 100 epochs using the AdamW optimizer (learning rate = 1 *×* 10^−4^), a batch size of 128, and cross-entropy loss with L1 (λ_1_ = 0.01) and L2 (λ=1 *×* 10^−4^) regularization.

#### 2.5.4 Gaussian Naive Bayes

A Gaussian Naive Bayes classifier was implemented using scikit-learn’s GaussianNB. Input features were standardized using parameters estimated from the training data and subsequently applied to the test data. The classifier modeled each input feature using a class-conditional Gaussian distribution and combined the resulting likelihoods under the Naive Bayes conditional-independence assumption. The default variance smoothing parameter of 1 *×* 10^*−*9^ was used [72].

#### 2.5.5 Mixture of Experts (MoE)

The primary classification model employed a sparsely gated Mixture of Experts (MoE) architecture, implemented in PyTorch [10]. The MoE framework combines multiple specialized expert networks with a trainable gating network that learns to route each input sample to the most relevant experts [35, 52].

##### Gating Network

The gating mechanism computes a sparse distribution over experts for each input. Given an input vector *x*, the gating network produces logits using a learned weight matrix (*W*_*g*_):

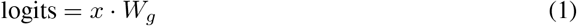

To encourage exploration during training, noisy gating was employed by adding tunable Gaussian noise scaled by a learned noise matrix (*W*_*n*_) and a Softplus activation:

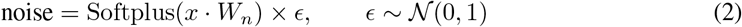

The noisy logits were passed through a softmax function, and only the top-*k* experts (*k* =4 out of 10 total experts) were selected for each input. The gating weights were then renormalized over the selected experts. This sparsity ensures computational efficiency because only a subset of experts is activated for each sample [15].

##### Expert Networks

To effectively capture the multimodal nature of synthetic riboswitch constructs, the MoE architecture employed a heterogeneous ensemble of 10 specialized expert networks: seven multilayer perceptron (MLP) modules dedicated to processing the 26 ViennaRNA thermodynamic and structural descriptors, and three multi-head self-attention sequence modules designed to model spatial base-pairing and positional dependencies across the one-hot encoded RNA sequence. Each MLP expert consisted of a two-layer feedforward network (26 → 64 → 2) with ReLU activation. Each sequence expert mapped the 4-dimensional nucleotide embeddings to a 64-dimensional latent space with learned 1D positional encodings, followed by a 4-head multi-head attention layer (`embed_dim=64`, `dropout=0.1`), average sequence pooling, and a linear projection to the two-class output space. A learned sparse gating network routed each input construct to the top-*k* (*k* = 4) most relevant experts per sample [10].

##### Sparse Dispatch

A sparse dispatcher routed input samples to their assigned experts based on the gating decisions, computed expert outputs independently, and combined them through a weighted sum using the gating weights.

##### Loss Function

The model was trained using cross-entropy loss [59] for binary classification. An auxiliary load-balancing loss was added to prevent expert collapse by encouraging balanced expert utilization. This loss was computed as the squared coefficient of variation (CV^2^) of both the expert importance (sum of gating weights) and expert load (number of assigned samples):

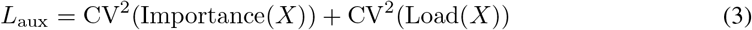

The total training loss was the sum of the cross-entropy loss and the auxiliary load-balancing loss.

##### Training Configuration

The model was trained for 100 epochs using the Adam optimizer with a learning rate of 1 × 10^*−*4^, a batch size of 128, and both L1 (*λ*_1_ = 0.01) and L2 (*λ*_2_ =5 × 10^*−*4^) regularization to control overfitting [47, 48]. L2 regularization was applied through weight decay in the optimizer, whereas L1 regularization was computed as the sum of the absolute values of all model parameters and added directly to the total loss.

### 2.6 Ablation Analysis

Ablation experiments were performed to evaluate the relative contribution of the numerical and sequence-derived inputs to classification performance. Two complementary analyses were conducted: numerical-feature ablation and input-source ablation. The same training and test partitions were used across experiments.

#### 2.6.1 Individual Numerical Feature Ablation

Individual-feature ablation were first performed to assess sensitivity of each model to the 26 numerical features. For each experiment, one numerical feature was independently set to zero while the remaining 25 features and 840-dimensional one-hot encoded sequence representation were retained. After each experiment, the ablation feature was restored before next numerical feature was evaluated. Only one numerical feature was set to zero in each individual-feature experiment and models were retrained and evaluated independently for every ablation condition.

Because the Decision Tree showed the greatest sensitivity to individual feature ablation, its results were used to rank the 26 numerical features. Features producing the largest decrease in Decision Tree accuracy were ranked first. This ranking was subsequently used to determine the order of the cumulative ablation experiment. Detailed individual-feature results are provided in the Supplementary Information.

#### 2.6.2 Cumulative Numerical Feature Ablation

Using the Decision Tree-derived ranking, numerical features were progressively set to zero to evaluate model performance as increasing amounts of numerical information were ablated. Unlike the individual-feature analysis, previously zeroed features remained zero as additional features were ablated. The native 840-dimensional one-hot encoded sequence representation was retained unchanged throughout the experiment.

The analysis ranged from the full-input condition, with no numerical features zeroed, to a sequence-only condition in which all 26 numerical features were zeroed. Each condition was independently trained and evaluated using the Decision Tree, Random Forest, Dense Neural Network, Gaussian Naive Bayes, and Mixture of Experts models.

#### 2.6.3 Numerical and Sequence Input Ablation

An input-source ablation experiment was performed to determine whether the numerical features alone were sufficient for classification. In the numerical-only condition, all 26 numerical features were retained, while the native sequence of each sample was replaced with the same dummy 210-nucleotide sequence composed entirely of adenines (A). The dummy sequence was one-hot encoded in the same manner as the native sequences, producing an identical 840-dimensional sequence representation for all samples and therefore removing sample-specific sequence information.

All five models were evaluated under three input conditions: full input (numerical features + native sequence), sequence only (all 26 numerical features set to zero), and numerical only (all 26 numerical features + constant poly-A sequence).

### 2.7 Evaluation Protocol

The dataset was split into training/validation (80%) and test (20%) partitions using a fixed random seed (random_state = 42). Within the training partition, 5-fold cross-validation with shuffled splits was employed to assess model generalization. Performance was measured using accuracy, F1 score, precision, and recall for classification.

## 3 Results

### 3.1 Machine Learning Comparison on CVB3

We evaluated the MoE classifier and other models’ performance on the CVB3 dataset.

#### 3.1.1 Mixture of Experts (MoE) Performance

The Mixture of Experts classifier demonstrated rapid convergence during training, with both training and validation loss decreasing sharply within the first 10 epochs before stabilizing at low values for the remainder of training (Figure 3). The close agreement between the training and validation loss curves indicates stable optimization, with no evidence of divergence or unstable learning. Similarly, training and validation accuracy increased rapidly and reached 100% within the first several epochs (Figure 4). The overlapping accuracy curves indicate consistent performance across the 5-fold cross-validation process.

**Figure 3:**
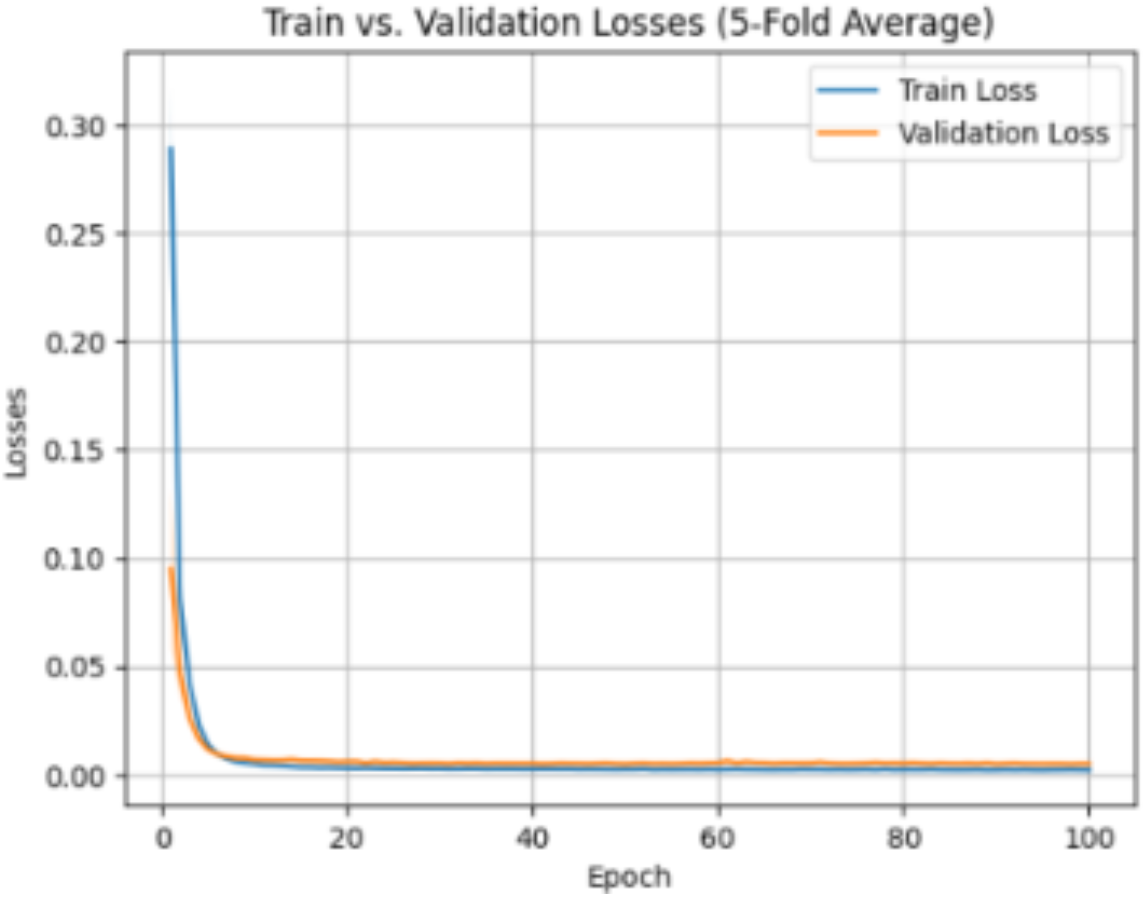
Training and validation loss of the MoE classifier during 5-fold cross-validation.

**Figure 4:**
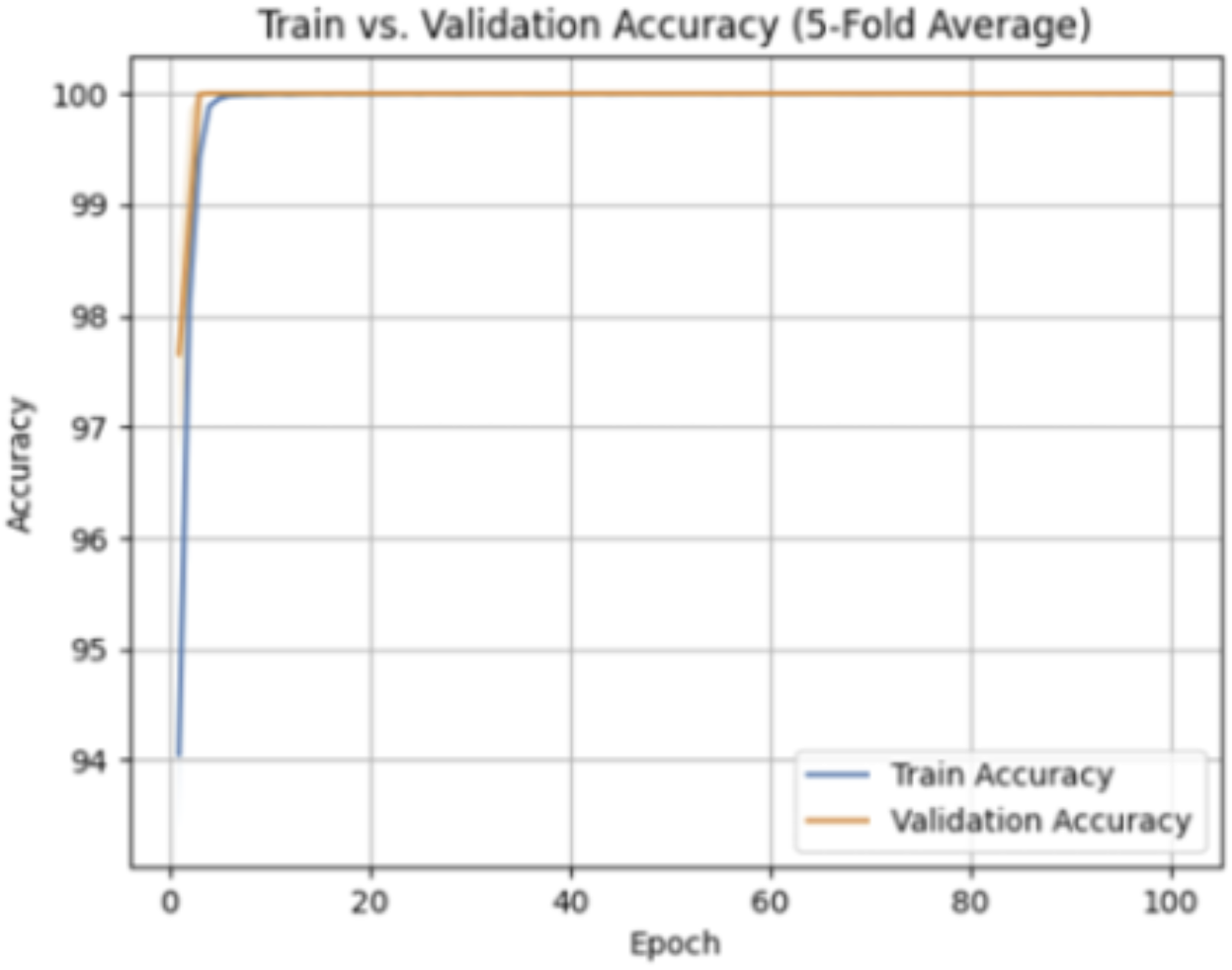
Training and validation accuracy of the MoE classifier during 5-fold cross-validation.

The cross-validation results demonstrated stable model performance across all folds (Table 2). Validation accuracy, precision, and recall reached 100% in every fold, while validation loss ranged from 0.0037 to 0.0073. The auxiliary loss remained low across folds, indicating stable expert utilization during training.

**Table 2:** Fold-by-Fold Validation Performance of the MoE Classifier.

| Fold | Accuracy (%) | Loss | Auxiliary Loss | Precision | Recall |
| --- | --- | --- | --- | --- | --- |
| 1 | 100 | 0.00379264 | 0.00379256 | 1 | 1 |
| 2 | 100 | 0.00533775 | 0.00533772 | 1 | 1 |
| 3 | 100 | 0.0037569 | 0.00375682 | 1 | 1 |
| 4 | 100 | 0.0051181 | 0.00511795 | 1 | 1 |
| 5 | 100 | 0.0073167 | 0.00731668 | 1 | 1 |

Evaluation on the independent test set produced identical classification performance (Figure 5, and Table 3). The confusion matrix showed that 5,497 correctly classified negative samples and 5,508 correctly classified positive samples, with no false positives or false negatives. Overall, the model was able to achieve 100% accuracy, precision, recall, and F1 score on a held-out test set. Although these results indicate excellent predictive performance, the uniformly perfect classification across training, validation, and test datasets motivated further investigation into whether the model was learning biologically meaningful patterns or exploiting dataset-specific characteristics.

**Table 3:**
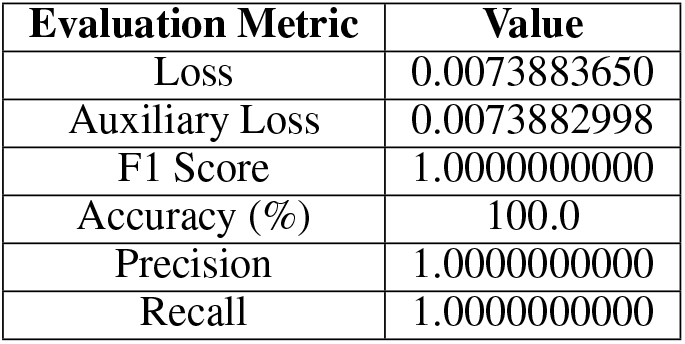
Performance Summary of the MoE Classifier on the Independent Test Set.

**Figure 5:**
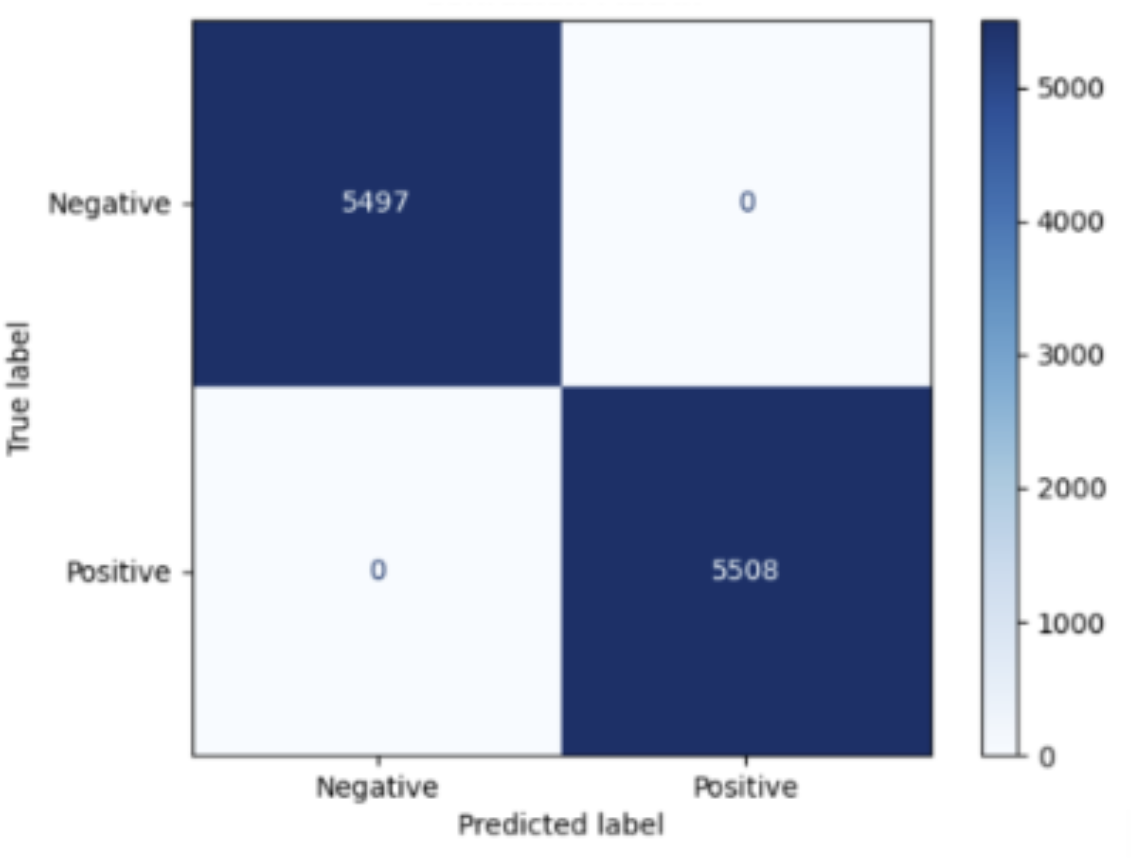
Confusion matrix of the MoE classifier on the independent test set.

#### 3.1.2 Dense Neural Network Performance

The Dense Neural Network was evaluated using the same 5-fold cross-validation setup as the MoE classifier. Using optimal regularization setting (L1 = 0.01, L2 = 5 * 10-4),the model converged rapidly, with both training and validation accuracy approaching 100% within the first several epochs (Figure 6A). Similarly, both the training and validation loss decreased rapidly and remained close to zero for the remainder of training (Figure 6B), indicating stable convergence under the selected regularization parameters.

**Figure 6:**
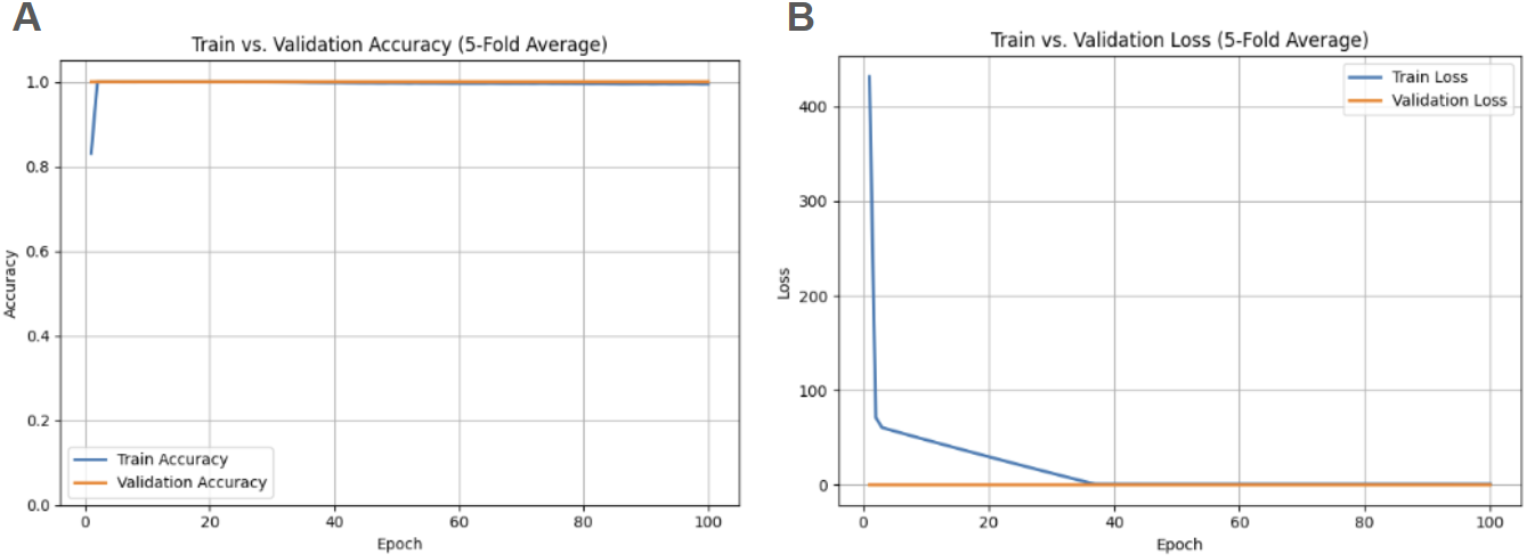
Training dynamics of the Dense Neural Network during 5-fold cross-validation using L1 = 0.01 and L2 = 5 × 10^*−*4^. (A) Mean training and validation accuracy across the five cross-validation folds. (B) Mean training and validation loss across the five cross-validation folds. The model converged rapidly and maintained consistently high validation performance throughout training.

Table 4 summarizes the classification performance of the Dense Neural Network classifier. The model achieved perfect classification performance across all evaluation metrics during both cross-validation and independent testing. The average 5-fold cross-validation accuracy, precision, recall, and F1-score were all 1.000, and the independent test set produced identical values with a test loss of 0.028. No false-positive or false-negative predictions were observed in the independent test confusion matrix (Figure 7). Similar performance was observed in the MoE classifier, suggesting that the dataset contains highly informative class-associated features. While the Dense Neural Network classifier demonstrated excellent predictive performance, the uniformly perfect results indicate that feature informativeness and dataset structure, rather than model architecture alone, likely contributed substantially to classification performance.

**Table 4:** Dense Neural Network Classification Performance (L1 = 0.01, L2 = 5 *×* 10^*−*4^)

| Evaluation | Accuracy | Precision | Recall | F1-score | Loss |
| --- | --- | --- | --- | --- | --- |
| 5-Fold Cross-Validation Average | 1.000 | 1.000 | 1.000 | 1.000 | — |
| Independent Test Set | 1.000 | 1.000 | 1.000 | 1.000 | 0.028 |

**Figure 7:**
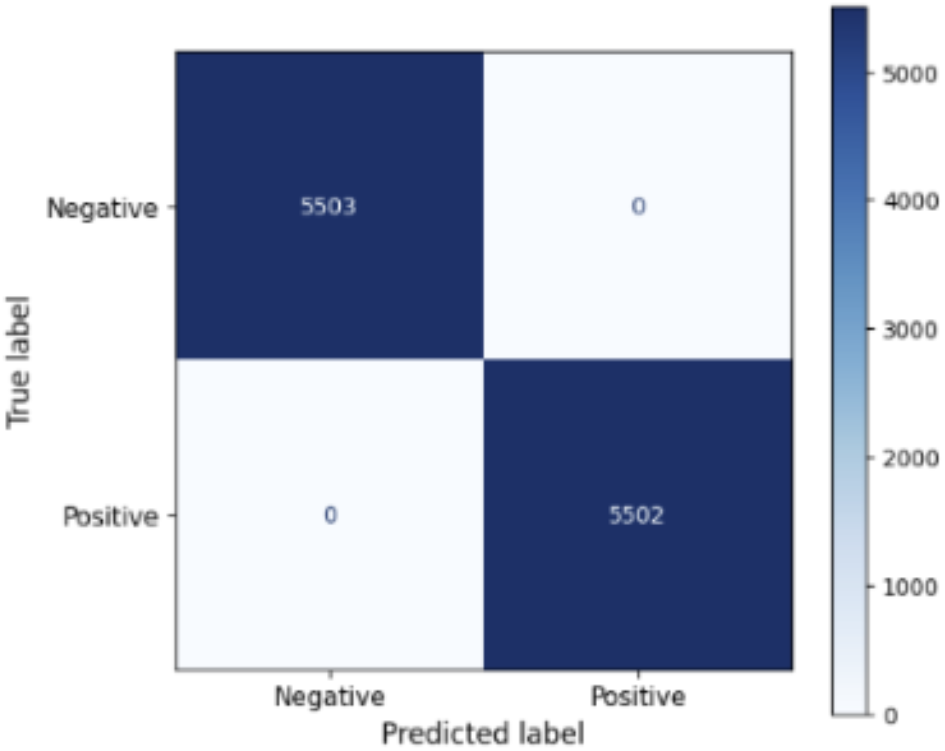
Confusion matrix of the Dense Neural Network on the independent test set.

#### 3.1.3 Decision Tree Performance

To further investigate the source of the consistently high performance observed across neural network models, a Decision Tree classifier was trained as an interpretable baseline.

The Decision Tree achieved a high performance across both 5-fold cross-validation and independent test datasets (Table 5). The average cross-validation accuracy was 0.9886 ± 0.0025, while the model achieved a test accuracy of 0.9926, with corresponding precision, recall, and F1-score values of 0.9963 ± 0.0015, 0.9808 ± 0.0065, and 0.9885 ± 0.0026, respectively. Evaluation on the independent test set had an accuracy of 0.9926, with a precision of 0.9991, recall of 0.9862, and an F1-score of 0.9926. Overall, these results indicate that the Decision Tree was able to achieve predictive performance comparable to more complex neural network architectures.

**Table 5:** Decision Tree Classifier Performance.

| Evaluation | Accuracy | Precision | Recall | F1-score |
| --- | --- | --- | --- | --- |
| Fold 1 Val | 0.9863 | 0.9968 | 0.9757 | 0.9861 |
| Fold 2 Val | 0.9894 | 0.9952 | 0.9836 | 0.9894 |
| Fold 3 Val | 0.9883 | 0.9981 | 0.9784 | 0.9882 |
| Fold 4 Val | 0.9928 | 0.9941 | 0.9916 | 0.9928 |
| Fold 5 Val | 0.9913 | 0.9972 | 0.9748 | 0.9859 |
| **CV Mean ± Std** | **0.9886 ± 0.0025** | **0.9963 ± 0.0015** | **0.9808 ± 0.0065** | **0.9885 ± 0.0026** |
| **Independent Test** | **0.9926** | **0.9991** | **0.9862** | **0.9926** |

The independent test confusion matrix (Figure 8) demonstrates that the classifier was able to correctly identify 5,498 negative samples and 5,426 positive samples while producing only 5 false positives and 76 false negative predictions. The small number of classification errors further supports the model’s strong generalization performance on the previously unseen data.

**Figure 8:**
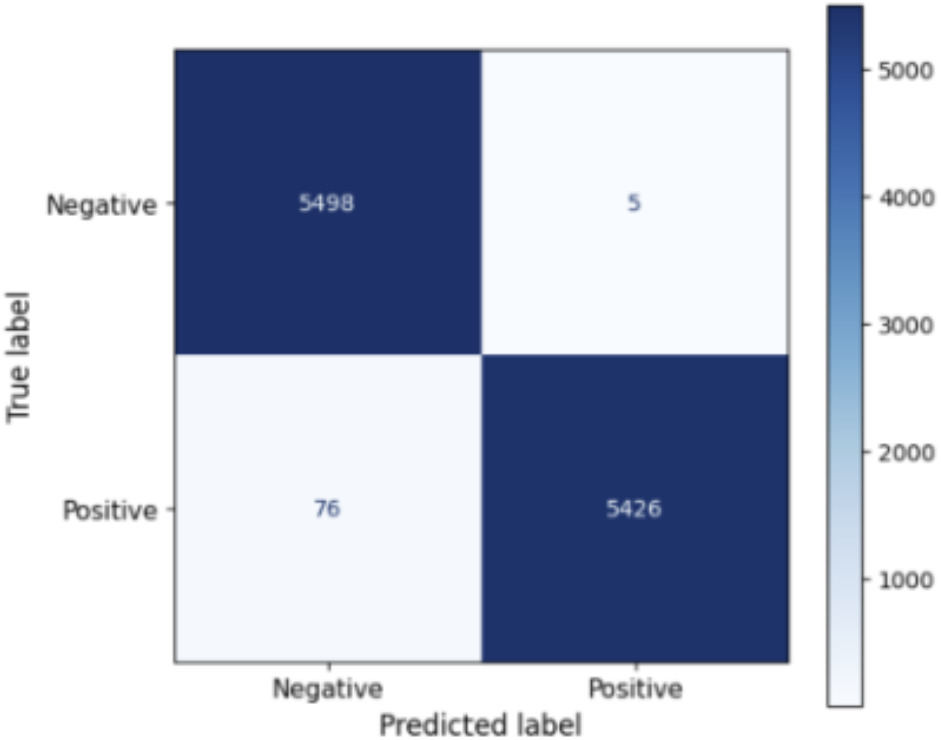
Confusion matrix of the Decision Tree evaluated on the independent test set.

To better understand which variables are driving the model’s predictions, feature importance scores were extracted from the final trained Decision Tree (Figure 9). Among all the features, base_636 accounted for approximately 86.5% of the total feature importance, which exceeds every other feature. The next most influential feature is reporter_single_strandedness, which contributed only a small fraction of the overall importance. The highly skewed distribution indicates that the classifier relied primarily on a single sequence-derived feature, rather than integrating information across many biological variables.

**Figure 9:**
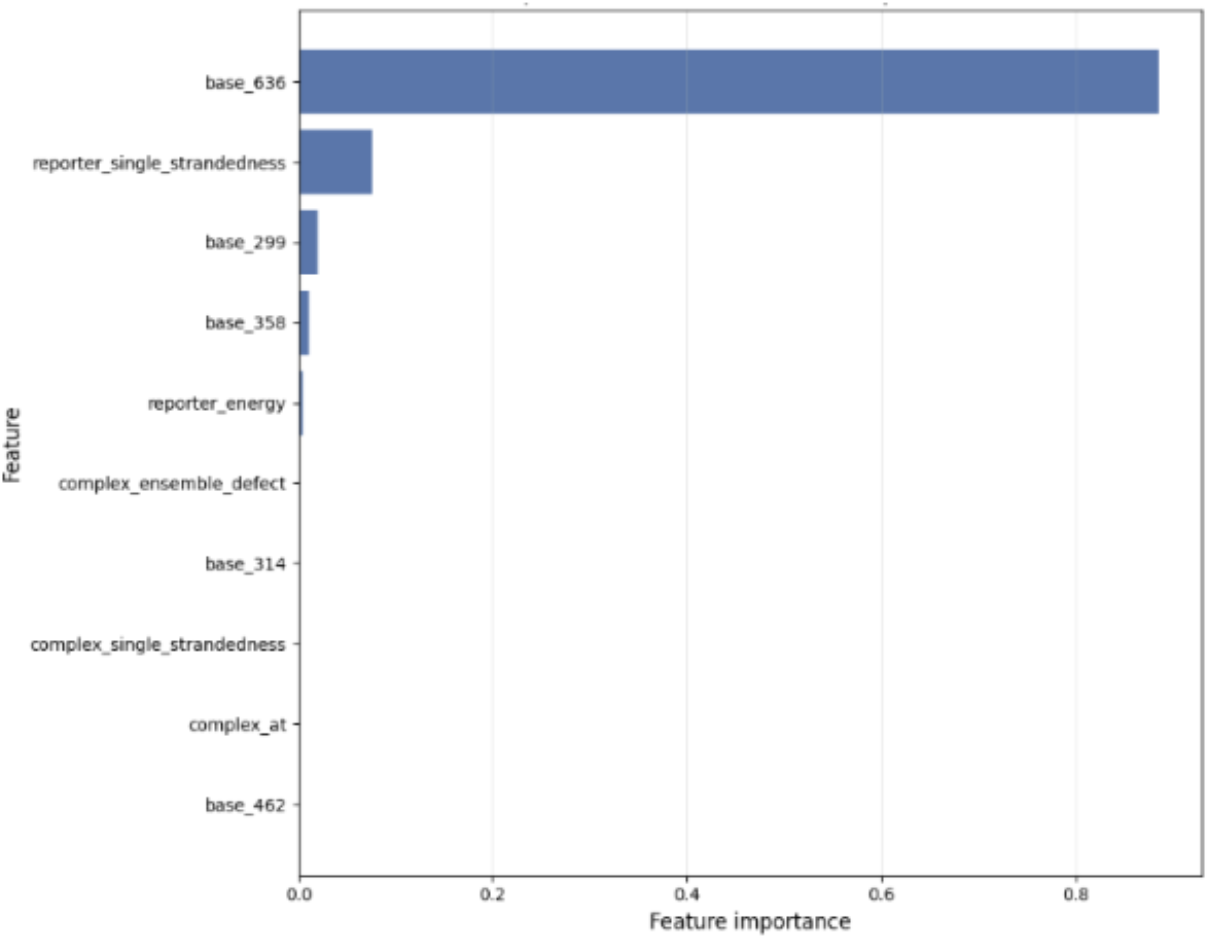
Top 10 Decision Tree feature importances.

The dominance of base_636 suggests that the dataset contained one or more highly discriminative predictors. These findings motivated a more detailed investigation of potential feature leakage.

#### 3.1.4 Comparative Performance Summary

All five machine learning models were evaluated on the independent test set (11,005 constructs) using the standardized 866-dimensional feature representation comprising 26 thermodynamic and structural descriptors alongside 840 sequence-derived one-hot features. Across these models, four architectures (the Mixture of Experts classifier, Dense Neural Network, Random Forest, and Gaussian Naive Bayes) achieved 100.00% test accuracy and a test F1-score of 1.000, correctly classifying every held-out construct without a single false positive or false negative prediction. The baseline Decision Tree achieved 99.26% test accuracy and an F1-score of 0.9926, producing 81 classification errors (5 false positives and 76 false negatives). As detailed in Section 3.1.1, the Decision Tree predictions were overwhelmingly driven by a single one-hot sequence position, `base_636`, which accounted for 86.5% of total feature importance. Table 6 summarizes the comparative performance metrics and structural specifications across all five models.

**Table 6:** Comparative Performance Summary Across Evaluated Classification Models.

| Method | Test Accuracy | Test F1 | Training Time | Configuration / Parameters |
| --- | --- | --- | --- | --- |
| MoE Classifier | 100.00% | 1.000 | 834.62 s | 10 specialized experts (7 MLP, 3 MHA); $h = 64$; $k = 4$; test loss = 0.0074 |
| Dense NN Classifier | 100.00% | 1.000 | 41.18 s | 4-layer MLP; LayerNorm; LeakyReLU; Dropout ($p = 0.3$); 100 epochs; test loss = 0.0283 |
| Random Forest | 100.00% | 1.000 | 4.52 s | 300 decision trees; max_features = 'sqrt' |
| Gaussian Naive Bayes | 100.00% | 1.000 | 0.05 s | Standardized input features; variance smoothing = $1 \times 10^{-9}$ |
| Decision Tree | 99.26% | 0.9926 | 0.15 s | Depth = 24; 277 leaves; max_features = 'sqrt' |

Despite the parity in test accuracy across the four leading architectures, the Mixture of Experts model offers distinct advantages. When comparing the two gradient-optimized neural network frameworks, the MoE classifier achieved a lower cross-entropy test loss (0.0074) than the Dense Neural Network (0.0283), reflecting sharper confidence calibration on unseen constructs. More importantly, the MoE design provides modular inductive biases by structurally separating scalar thermodynamic inputs (processed by MLP experts) from nucleotide sequences (processed by multi-head self-attention experts). The learned top-*k* gating mechanism dynamically routes samples to relevant subnetworks, providing high representational capacity while evaluating only a sparse subset of parameters per construct.

Although all five models demonstrated strong predictive performance on the benchmark, the emergence of near-perfect classification across diverse model families motivated a deeper investigation into whether the learned decision boundaries capture authentic physical RNA folding rules or exploit dataset-level structural artifacts.

### 3.2 Ablation Studies

#### 3.2.1 Cumulative Numerical-Feature Ablation

To evaluate how progressively reducing numerical information affected model performance, the 26 numerical features were cumulatively set to zero according to the Decision Tree sensitivity ranking while the native sequence representation was retained. Figure 10 shows the change in test accuracy as additional numerical features were ablated, while Figure 11 summarizes the same results across all five models.

**Figure 10:**
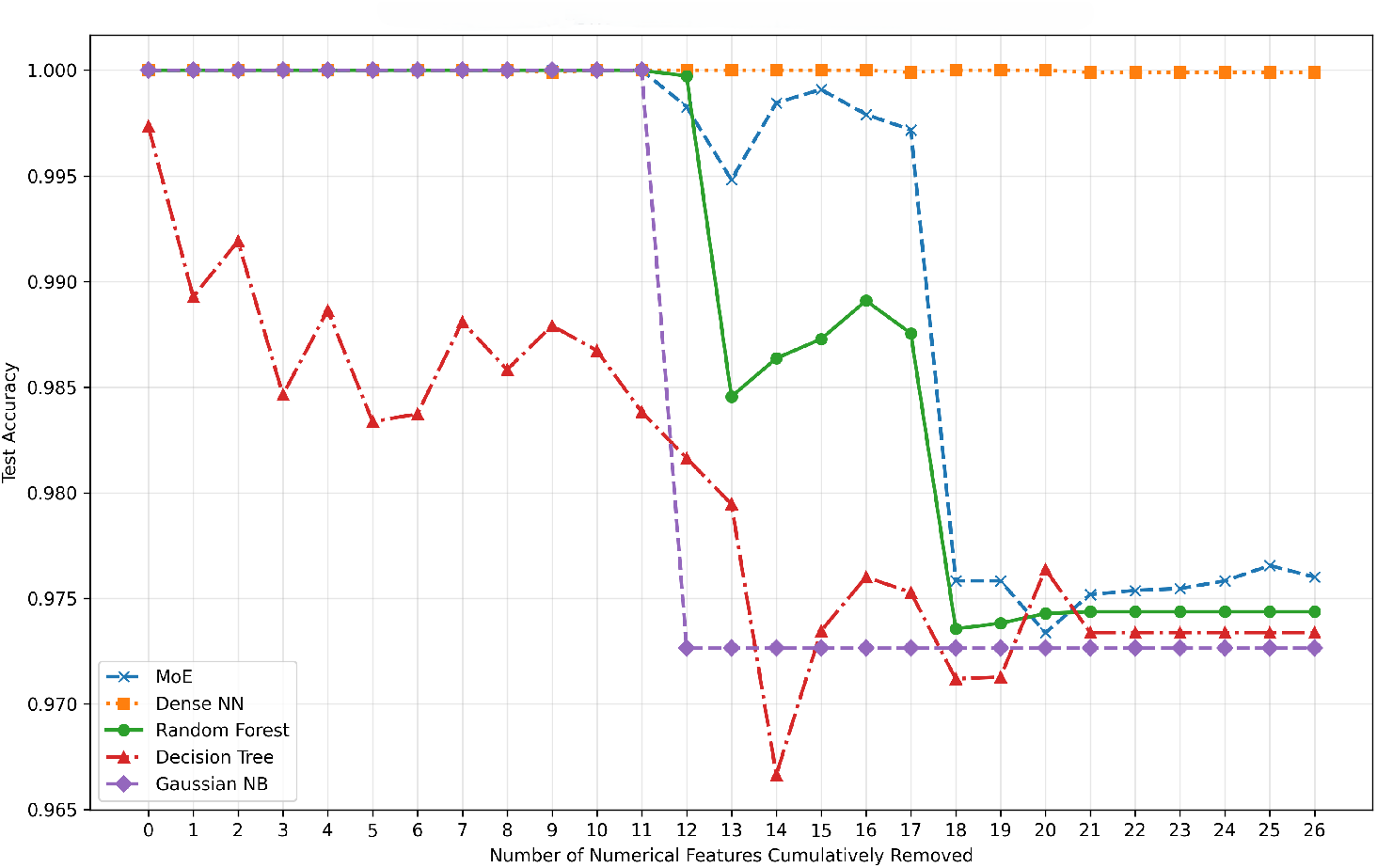
Cumulative Numerical Feature Ablation Across Five Models.

**Figure 11:**
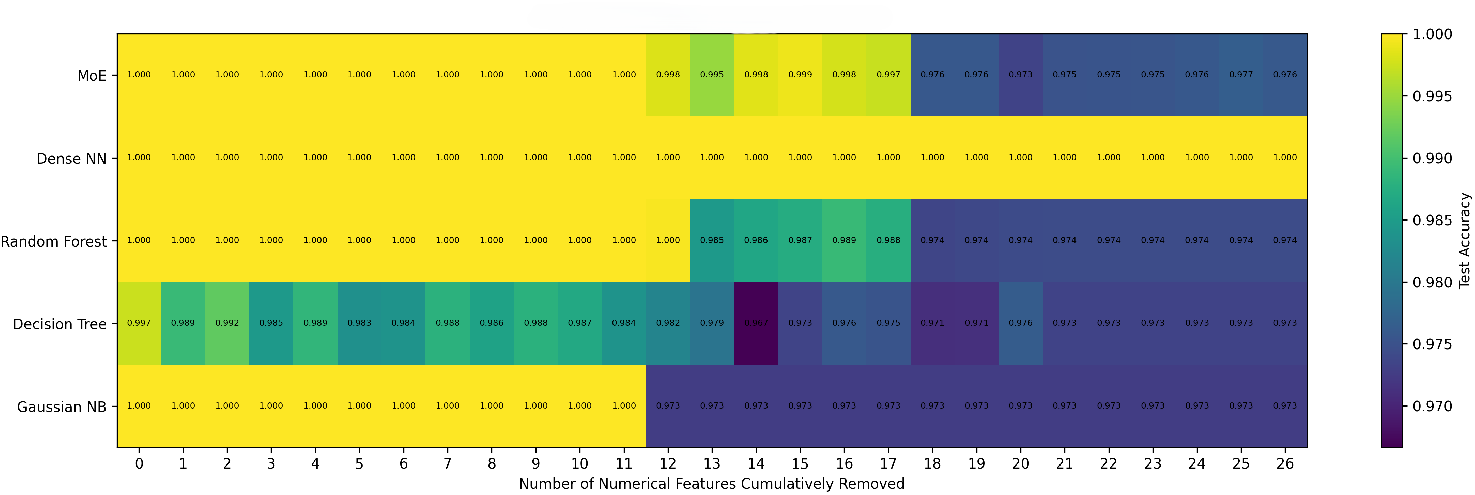
Heatmap of test accuracy across cumulative numerical-feature ablation conditions for all five classification models.

The cumulative ablation analysis revealed distinct levels of sensitivity across the five models. The Dense Neural Network maintained an accuracy of 1.000 across all ablation conditions. MoE also remained at 1.000 through the first 11 numerical features zeroed, after which performance declined modestly, reaching 0.976 when all 26 numerical features were zeroed. Random Forest showed a similar pattern, with accuracy decreasing to approximately 0.974 under the sequence-only condition.

Gaussian Naive Bayes remained at 1.000 through the first 11 ablations, then decreased to approximately 0.973 and remained stable. The Decision Tree was the most sensitive model, with accuracy varying between approximately 0.967 and 0.997 across conditions. Overall, all five models retained high predictive accuracy even when all 26 numerical features were zeroed, indicating that the retained sequence representation alone contained substantial class-discriminative information.

#### 3.2.2 Numerical Feature and Sequence Input Ablation

To determine whether the numerical and sequence inputs were independently sufficient for classification, the five models were evaluated under full-input, sequence-only, and numerical-only conditions. In the numerical-only condition, all 26 numerical features were retained while every sample received the same dummy 210-nucleotide sequence composed entirely of adenines. All five models achieved an accuracy of 1.000 in this condition. Under the sequence-only condition, accuracy ranged from 0.973 to 1.000 across the five models (Figure 12).

**Figure 12:**
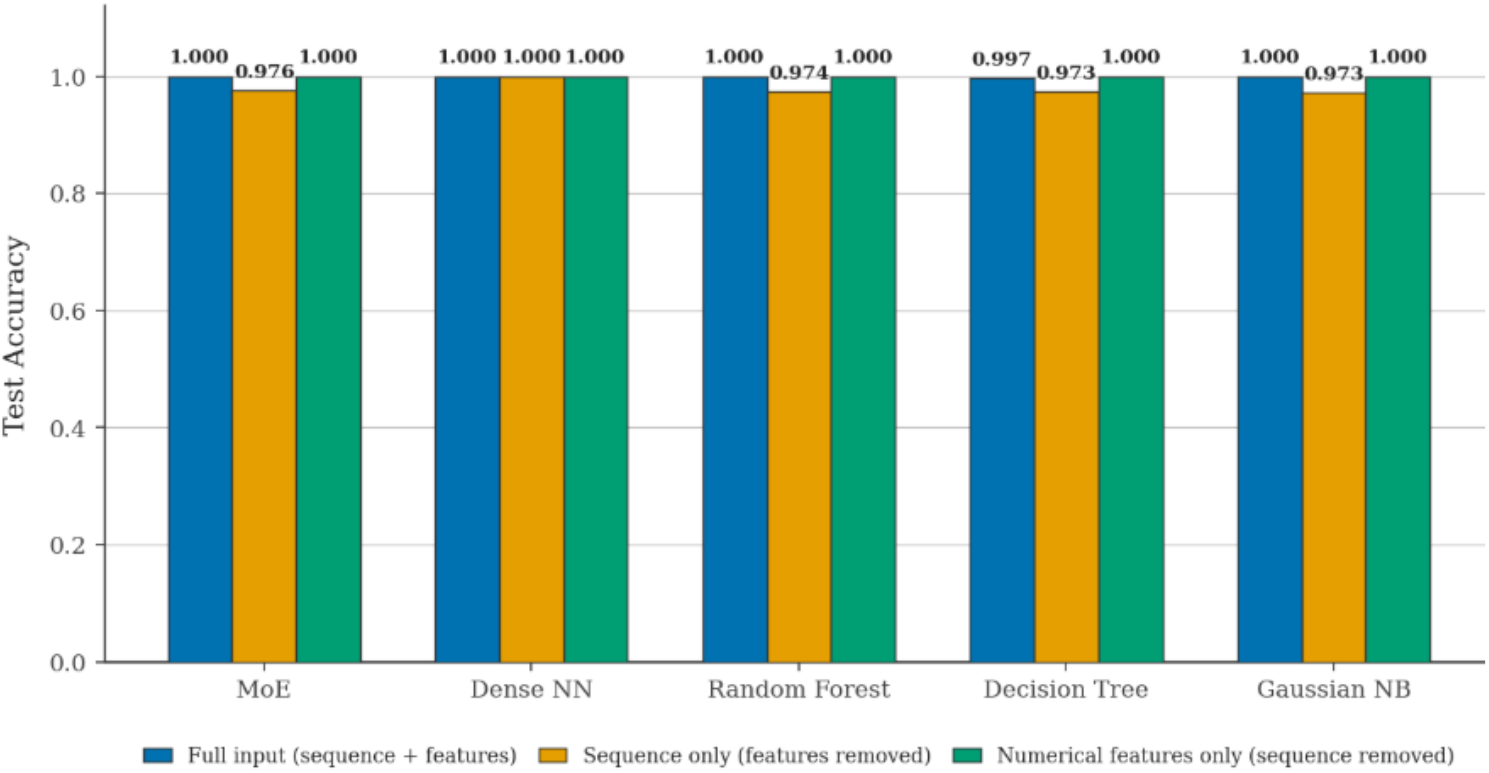
Model Performance Under Full-Input, Sequence-Only, and Numerical-Only Conditions

Although the Dense Neural Network achieved high accuracy across all ablation conditions, the MoE architecture was retained as the primary model because its sparse gating mechanism selectively activates specialized subnetworks for each input modality, offering greater representational capacity and modular inductive bias. This design can provide greater model capacity without requiring all model parameters to be activated for every sample and offers a potential computational advantage over a fully dense architecture while maintaining high performance.

### 3.3 Performance of the MoE Classifier in a Mixed-Virus Context

To further assess the model’s performance, we evaluated the MoE classifier on a mixed sample containing CVB3 and MS2. In practice, multiple viruses may coexist in the same environment. A model capable of simultaneous detection of multiple viruses could support multiplexed analysis and potentially reduce the resources required for separate virus detection. This capability could help extend the framework to practical environmental monitoring and pathogen detection in diagnostic applications [75, 74]. From a modeling perspective, the framework may also provide a more flexible approach than training separate classifiers for each virus.

The MoE classifier was trained and tested on a data sample consisting of CVB3 and MS2 at a salt concentration of 0.01 M, with 27,512 samples for each virus. The data were divided into training and testing sets at an 80:20 ratio. The target sequence of MS2 was derived from the RNA polymerase locus (positions 1761-3398 bp) obtained from NCBI [73]. The model was trained for 100 epochs with L1 (*λ*_1_ =1 *×* 10^*−*3^) and L2 (*λ*_2_ =1 *×* 10^*−*3^) regularization.

Figure 13 illustrates the training and validation results. The results showed steady training convergence, where both the training and validation losses approached zero by epoch 40, and accuracy rapidly approached 100%. The classifier also demonstrated strong performance on the test set (Figure 14 and Table 7). The confusion matrix showed that all samples were classified correctly with no false positives or false negatives. The test loss was 0.0698, and the accuracy, precision, recall, and F1 score were all 100%. These results demonstrate perfect classification on the evaluated mixed-virus test set and suggest the framework could potentially be extended to viruses other than CVB3 while maintaining high classification performance. Overall, the findings highlight the classifier’s potential for flexible multi-virus detection, while further evaluation across additional viruses, salt concentrations, and virus combinations is needed to assess the generalizability and scalability of the framework.

**Table 7:** Performance Summary of the MoE Classifier on the Independent Test Set containing CVB3 and MS2 Samples.

| Evaluation Metric | Value |
| --- | --- |
| Loss | 0.0698288908 |
| Auxiliary Loss | 0.0172180008 |
| F1 Score | 1.0000000000 |
| Accuracy (%) | 100.0 |
| Precision | 1.0000000000 |
| Recall | 1.0000000000 |

**Figure 13:**
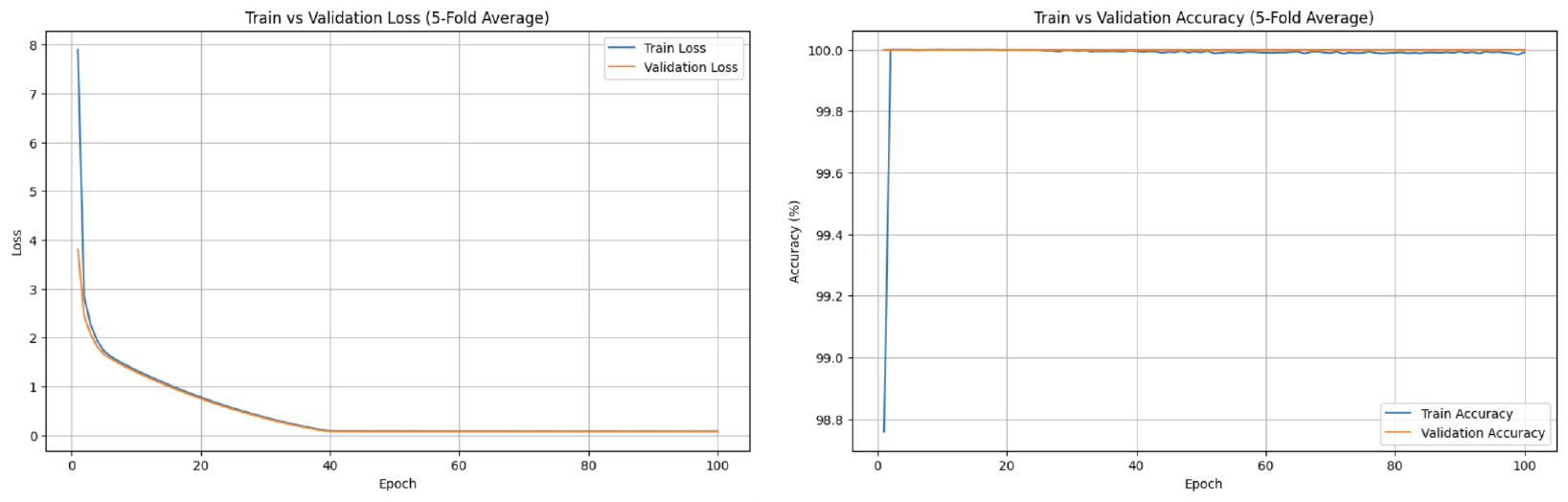
Training and validation losses (left) and accuracies (right) of the MoE classifier during 5-fold cross-validation for CVB3 and MS2 classification

**Figure 14:**
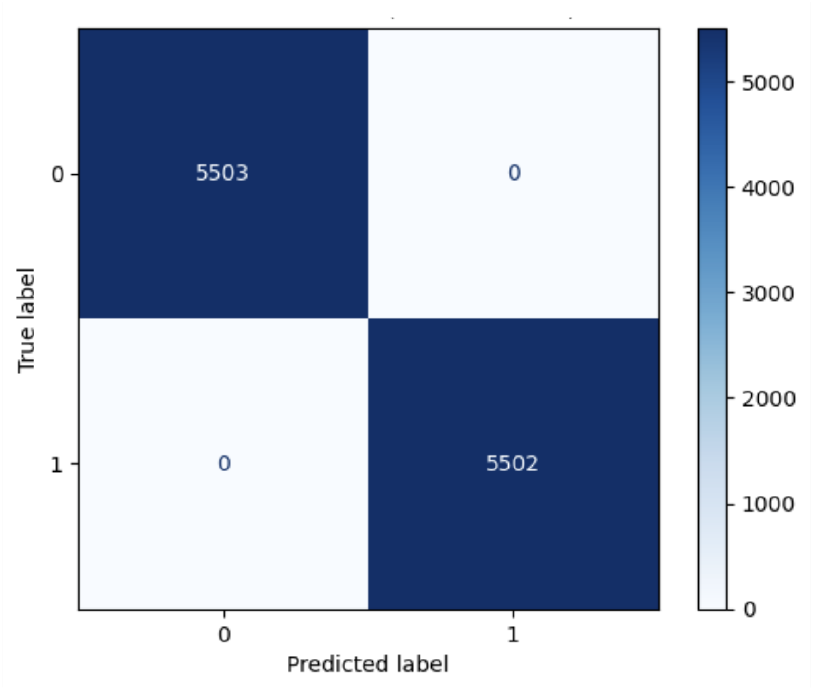
Confusion matrix of the MoE classifier on the independent test set containing CVB3 and MS2 samples.

### 3.4 Sequence-Level Feature Analysis

To further analyze the balanced riboswitch dataset, we examined the distributions of key structural and thermodynamic features for positive and negative constructs. Among these features exhibiting within-class variation, complex_energy and complex_single_strandedness demonstrated the greatest separation between classes (Figure 15). Positive constructs showed a more negative mean complex_energy (−78.3) than negative constructs (−44.1), indicating lower probability of nucleotides remaining unbound within the complex. Although distributions overlapped, both features showed clear shifts between classes, with separation scores of 1.78 and 1.67, respectively. The full set of results are given in Supplementary Information A.1.

**Figure 15:**
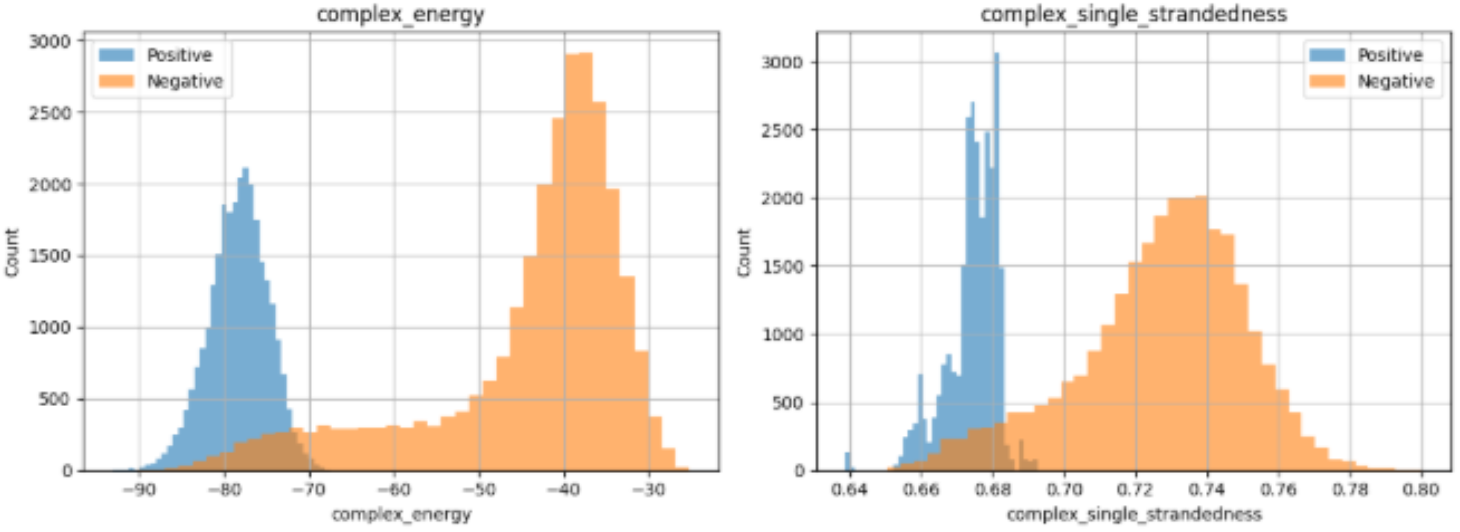
Distribution of `complex_energy` and `complex_single_strandedness`.

In contrast, reporter_energy, reporter_ensemble_defect, and reporter_single_strandedness exhibited even larger apparent class separation, but assumed a single constant value within each class. Since these features lacked within-class variability, they were not considered representative of genuine biological variation and were excluded from further structural interpretation.

### 3.5 K-mer Analysis

Windowed 3-mer analysis showed stronger positional variation in the Coxsackievirus B3 (CVB3) target sequences than in the MS2 sequences, indicating greater variation in local sequence composition across the CVB3 target region (Figure 16). In contrast, the MS2 target and trigger sequences showed more uniform k-mer profiles across window positions. Additional windowed k-mer analyses and supporting statistics are provided in Supplementary Information A.2.

**Figure 16:**
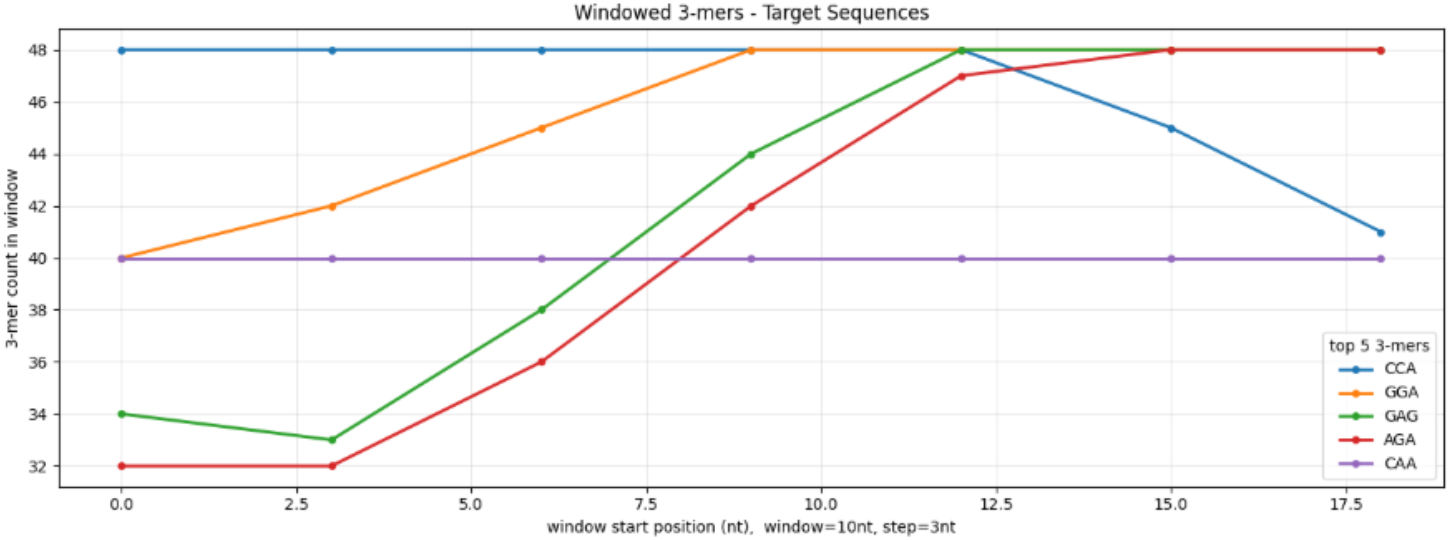
Windowed 3-mer counts for CVB3 target sequences.

**Figure 17:**
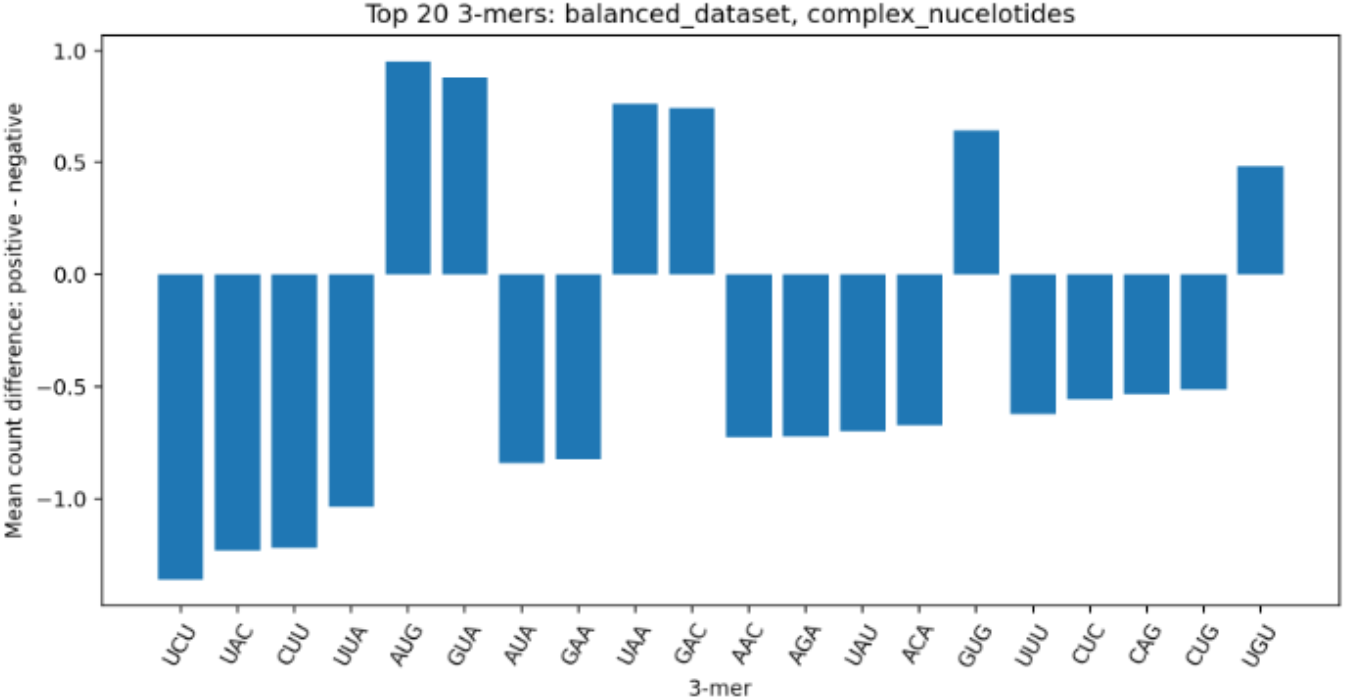
Top 20 3-mers by mean-count difference between positive and negative constructs, computed on the whole riboswitch complex sequence (balanced dataset, 27,512 positive and 27,512 negative).

## 4 Discussion

### 4.1 Biological Interpretation and Dataset-Specific Signals

The models achieved high classification performance when trained on the balanced riboswitch dataset; however, the feature analyses indicate that this performance should be interpreted cautiously. Positive constructs showed more negative complex_energy and lower complex_single_strandedness than negative constructs, suggesting that full-complex folding stability and accessibility differ between correctly paired and mismatched trigger-target constructs. These features are biologically plausible descriptors because RNA folding stability and nucleotide accessibility are expected to influence trigger-target annealing and riboswitch activation.

At the same time, several dataset-specific signals were identified. The Decision Tree relied heavily on base_636, indicating that a fixed-position sequence feature carried strong class information. Reporter-derived features, including reporter_energy, reporter_ensemble_defect, and reporter_single_strandedness, also showed class-specific non-overlapping values and were treated as leakage artifacts rather than biological signal. These findings suggest that the near-perfect performance of the MoE and Dense Neural Network classifiers cannot be interpreted solely as evidence of generalizable riboswitch-function prediction. Future models should therefore remove or recompute reporter-derived features, evaluate performance after masking fixed-position sequence shortcuts, and test generalization on datasets with balanced design-generation metadata and experimentally validated riboswitch activity.

### 4.2 Sequence-Level Insights from K-mer Analysis

The k-mer analysis provided an interpretable view of local sequence composition across the riboswitch datasets. Coxsackievirus B3 showed stronger positional variation in frequent 3-mers than MS2, suggesting greater local motif variation across the CVB3 target region. The size-matched MS2 subsampling analysis indicated that this difference was not explained by sample size alone.

Positive-versus-negative k-mer comparisons showed that target and trigger sequences alone had only minor class-associated differences. In contrast, the complete riboswitch complex showed substantially larger differences, suggesting that class-associated sequence-composition patterns are more pronounced in the full construct context. However, these results should be interpreted as descriptive sequence-composition differences rather than evidence that individual k-mers causally determine binding. Establishing binding relevance would require additional experimental validation or targeted folding and mutational analyses.

## 5 Conclusion and Future Work

Overall, this study demonstrates that integrating sequence-based and structural features can provide an effective framework for predicting riboswitch functions. Across the evaluated machine learning approaches, the Mixture of Experts model achieved the strongest predictive performance, demonstrating the value of combining complementary feature representations rather than relying on a single sequence or structural descriptor. These findings support the use of machine learning approaches for identifying patterns associated with riboswitch behavior and provide a computational framework for evaluating candidate riboswitch designs.

Feature-level analysis further provides insight into the sequence and structural characteristics associated with model predictions. K-mer analysis revealed that class associated sequence composition differences were substantially more pronounced when the complete riboswitch complex was considered than when target and trigger sequences were analyzed independently. Structural and thermodynamic descriptors provide complementary information related to RNA folding and trigger-target interactions. Together, these results suggest that riboswitch function is better characterized by considering multiple levels of sequence and structural information rather than individual sequence features in isolation.

Further work should validate the proposed framework using experimentally characterized riboswitch datasets and additional viral targets in order to determine whether sequence-composition patterns identified by the k-mer analysis have functional roles in riboswitch binding or activation. In addition, alternative RNA representations, including graph-based structural encoding and pretrained RNA language models could be investigated to improve predictive performance. Ultimately, integrating computational predictions with experimental validation could support more efficient identification and design of functional riboswitch candidates.

## 6 Dataset Availability

The datasets used in this study are available as follows:

**Balanced positive/negative CVB3 dataset used for model training and evaluation:**

https://drive.google.com/file/d/1mWpSsacMm3wD3uU8nXo1MqBrPOxN_AzQ/view?usp=drive_link

**Datasets used for kmer analysis**

**Green-Cherry-Cox-Data.csv (0.01) – CVB3:**

https://drive.google.com/file/d/11J4MeXKYyh9J1-WBdc4Z7CRnlArm_hlK/view?usp=sharing

**Green-LacZ-Cox-Data.csv (0.01) – CVB3:**

https://drive.google.com/file/d/1tSYaiCm20aAk6AY6Z4IsH-R2ytF-35BK/view?usp=sharing

**Green-MS2-Cherry-Data.csv (0.5) – MS2:**

https://drive.google.com/file/d/11cj-1AF5l4g2Gh6QIxemR1a3fVakblnk/view?usp=sharing

**Green-MS2-LacZ-Data.csv (0.5) – MS2:**

https://drive.google.com/file/d/1i-35Cv6KXzEnOaNtbwmK9HYRi4ys6FY4/view?usp=sharing

**Datasets used for training and evaluating mixed-virus classification model: Green-Cherry-Cox-Data.csv (0.01) – CVB3:**

https://drive.google.com/file/d/1SbzDZ4BCo4FX3Ii2x_C9V4VDXbM66T6Z/view?usp=sharing

**Green-MS2-Cherry-Data.csv (0.01) – MS2:**

https://drive.google.com/file/d/1yuLIq1pXcBWPJCymJpFwc901mkE3ZTd2/view?usp=sharing

## 7 Code Availability

**Original research and model-development code:**

https://github.com/Hungrycat112/RNA_water

**Standardized MoE implementation and Docker inference:**

https://github.com/Hungrycat112/riboswitch-moe

## 8 Contributions

JA provided mentorship and performed coding and data analysis. SH did analysis, visualizations and validation of results. BG and KC performed analysis, data processing and validation. HL and JJ coded and evaluated machine learning tools. JJ and SH wrote the container. AM and MT did analysis, validation of results and visualizations. WBA and LK supervised and mentored the project. All authors contributed to writing and editing the manuscript.

## 9 Acknowledgements

The authors gratefully acknowledge the SJSU College of Engineering’s Project Engineering Success and Summer Research Internship programs for their support. The authors thank the Department of Computer Science at SJSU for providing the high-performance computing resources that made the results in this paper possible.

## References

[1] Lohrmann, A., Farfan, J., Lohrmann, C., Kölbel, J. F., & Pettersson, F. (2023). Troubled waters: Estimating the role of the power sector in future water scarcity crises. Energy, 282, 128369.

[2] Farfan, J., & Lohrmann, A. (2023). Gone with the clouds: Estimating the electricity and water footprint of digital data services in Europe. Energy Conversion and Management, 290, 117215.

[3] Gerba, C. P., Betancourt, W. Q., & Kitajima, M. (2017). How much reduction of virus is needed for recycled water: A continuous changing need for assessment?. Water Research, 108, 25–31.

[4] Jonathan Auyong, Merylyn Negrete, William B. Andreopoulos, and Leila Khatib. Synthetic Riboswitch Sequences for CVB3 Detection Across Salt Concentrations. 2025 International Conference on Computational Science and Computational Intelligence (CSCI’25); American Council on Science and Education: Las Vegas, NV, December 3–5, 2025; In Proceedings of Springer Nature.

[5] Park, Soan, and Jeong Wook Lee. 2021. “Detection of Coronaviruses Using RNA Toehold Switch Sensors.” International Journal of Molecular Sciences 22 (4). 10.3390/ijms22041772.

[6] Pardee, Keith, Alexander A. Green, Tom Ferrante, D. Ewen Cameron, Ajay DaleyKeyser, Peng Yin, and James J. Collins. 2014. “Paper-Based Synthetic Gene Networks.” Cell 159 (4): 940–54.

[7] Green, Alexander A., Pamela A. Silver, James J. Collins, and Peng Yin. 2014. “Toehold Switches: De-Novo-Designed Regulators of Gene Expression.” Cell 159 (4): 925–39.

[8] Ogawa, Atsushi (ed.), Artificial Riboswitches: Methods and Protocols, Methods in Molecular Biology, Vol. 1111, DOI:10.1007/978-1-62703-755-6_1, Springer Science+Business Media New York 2014.

[9] L. Zeng, C. Otero, & L.S. Qi. (2025). Engineered RNA-based activation system for coronavirus sensing in live cells. Biosensors and Bioelectronics: X, 100040. 10.1016/j.bidere.2025.100040

[10] Hainian Audrey Long, “riboMoE: An Application of Mixture of Experts on Artificial Riboswitch Classification” (2025).Master’s Projects. 1510. DOI:10.31979/etd.yytx-j2ec

[11] “Riboswitches, RNA Regulation | Learn Science at Scitable,” [Online]. Available: https://www.nature.com/scitable/topicpage/riboswitches-a-common-rna-regulatory-element-14262702/

[12] B. L. Nicholson and K. A. White, “Functional long-range RNA–RNA interactions in positive-strand RNA viruses,” Nature Reviews Microbiology, vol. 12, no. 7, pp. 493–504, July 2014. [Online]. Available: https://www.nature.com/articles/nrmicro3288

[13] A. Lopez-del Rio, M. Martin, A. Perera-Lluna, and R. Saidi, “Effect of sequence padding on the performance of deep learning models in archaeal protein functional prediction,” Scientific Reports, vol. 10, no. 1, p. 14634, Sept. 2020. [Online]. Available: https://www.nature.com/articles/s41598-020-71450-8

[14] E. Rachid, “One hot encoding DNA,” Sept. 2020. [Online]. Available: https://elferachid.medium.com/one-hot-encoding-dna-92a1c29ba15a

[15] N. Shazeer, A. Mirhoseini, K. Maziarz, A. Davis, Q. Le, G. Hinton, and J. Dean, “Outrageously Large Neural Networks: The Sparsely-Gated Mixture-of-Experts Layer,” arXiv:1701.06538, Jan. 2017. [Online]. Available: http://arxiv.org/abs/1701.06538

[16] A. Vaswani, N. Shazeer, N. Parmar, J. Uszkoreit, L. Jones, A. N. Gomez, L. Kaiser, and I. Polosukhin, “Attention Is All You Need,” arXiv:1706.03762. [Online]. Available: http://arxiv.org/abs/1706.03762

[17] D. Garst, A. L. Edwards, and R. T. Batey, “Riboswitches: Structures and Mechanisms,” Cold Spring Harbor Perspectives in Biology, vol. 3, no. 6, p. a003533, June 2011. [Online]. Available: https://www.ncbi.nlm.nih.gov/pmc/articles/PMC3098680/

[18] C. Hammann and E. Westhof, “Searching genomes for ribozymes and riboswitches,” Genome Biology, vol. 8, no. 4, p. 210, 2007.

[19] S. Modrow, D. Falke, U. Truyen, and H. Schätzl, “Viruses with Single-Stranded, Positive-Sense RNA Genomes,” Molecular Virology, pp. 185–349, Aug. 2013. [Online]. Available: https://www.ncbi.nlm.nih.gov/pmc/articles/PMC7169642/

[20] J. Mamana, G. M. Humber, E. R. Espinal, S. Seo, N. Vollmuth, J. Sin, and B. J. Kim, “Coxsackievirus B3 infects and disrupts human induced-pluripotent stem cell derived brain-like endothelial cells,” Frontiers in Cellular and Infection Microbiology, vol. 13, p. 1171275, Apr. 2023. [Online]. Available: https://www.ncbi.nlm.nih.gov/pmc/articles/PMC10149843/

[21] “Coxsackievirus B3 - an overview | ScienceDirect Topics.” [Online]. Available: https://www.sciencedirect.com/topics/biochemistry-genetics-and-molecular-biology/coxsackievirus-b3

[22] K. D. Mena, C. P. Gerba, C. N. Haas, and J. B. Rose, “Risk Assessment of waterborne coxsackievirus,” Journal AWWA, vol. 95, no. 7, pp. 122–131, 2003. [Online]. Available: https://onlinelibrary.wiley.com/doi/abs/10.1002/j.1551-8833.2003.tb10413.x

[23] S. Duffy, “Why are RNA virus mutation rates so damn high?” PLoS Biology, vol. 16, no. 8, p. e3000003, Aug. 2018. [Online]. Available: https://www.ncbi.nlm.nih.gov/pmc/articles/PMC6107253/

[24] E. Domingo, C. García-Crespo, R. Lobo-Vega, and C. Perales, “Mutation Rates, Mutation Frequencies, and Proofreading-Repair Activities in RNA Virus Genetics,” Viruses, vol. 13, no. 9, p. 1882, Sept. 2021. [Online]. Available: https://www.ncbi.nlm.nih.gov/pmc/articles/PMC8473064/

[25] A. Cassedy, A. Parle-McDermott, and R. O’Kennedy, “Virus Detection: A Review of the Current and Emerging Molecular and Immunological Methods,” Frontiers in Molecular Biosciences, vol. 8, Apr. 2021. [Online]. Available: https://www.frontiersin.org/journals/molecular-biosciences/articles/10.3389/fmolb.2021.637559/full

[26] J. Zhang, Y. Fei, L. Sun, and Q. C. Zhang, “Advances and opportunities in RNA structure experimental determination and computational modeling,” Nature Methods, vol. 19, no. 10, pp. 1193–1207, Oct. 2022. [Online]. Available: https://www.nature.com/articles/s41592-022-01623-y

[27] H.-T. Yao, R. Lorenz, I. L. Hofacker, and P. F. Stadler, “Mono-valent salt corrections for RNA secondary structures in the ViennaRNA package,” Algorithms for Molecular Biology, vol. 18, no. 1, p. 8, July 2023. doi: 10.1186/s13015-023-00236-0.

[28] M. Zuker and P. Stiegler, “Optimal computer folding of large RNA sequences using thermodynamics and auxiliary information,” Nucleic Acids Research, vol. 9, no. 1, pp. 133–148, Jan. 1981.

[29] R. Lorenz, S. H. Bernhart, C. Höner zu Siederdissen, H. Tafer, C. Flamm, P. F. Stadler, and I. L. Hofacker, “ViennaRNA Package 2.0,” Algorithms for Molecular Biology, vol. 6, no. 1, p. 26, Nov. 2011. doi: 10.1186/1748-7188-6-26.

[30] “Vienna RNA web servers.” [Online]. Available: http://rna.tbi.univie.ac.at/RNAWebSuite/help.html

[31] J. S. McCaskill, “The equilibrium partition function and base pair binding probabilities for RNA secondary structure,” Biopolymers, vol. 29, no. 6–7, pp. 1105–1119, 1990.

[32] S. Wuchty, W. Fontana, I. L. Hofacker, and P. Schuster, “Complete suboptimal folding of RNA and the stability of secondary structures,” Biopolymers, vol. 49, no. 2, pp. 145–165, Feb. 1999.

[33] A. Shapiro and K. Zhang, “Comparing multiple RNA secondary structures using tree comparisons,” Bioinformatics, vol. 6, no. 4, pp. 309–318, 1990.

[34] T. Einert and R. Netz, “Theory for RNA Folding, Stretching, and Melting Including Loops and Salt,” Biophysical Journal, vol. 100, no. 11, pp. 2745–2753, June 2011.

[35] R. A. Jacobs, M. I. Jordan, S. J. Nowlan, and G. E. Hinton, “Adaptive Mixtures of Local Experts,” Neural Computation, vol. 3, no. 1, pp. 79–87, Mar. 1991.

[36] “Mixture of Experts Explained,” Feb. 2025. [Online]. Available: https://huggingface.co/blog/moe

[37] “deepseek-ai/DeepSeek-MoE,” Apr. 2025. [Online]. Available: https://github.com/deepseek-ai/DeepSeek-MoE

[38] DeepSeek-AI et al., “DeepSeek-V2: A Strong, Economical, and Efficient Mixture-of-Experts Language Model,” arXiv:2405.04434, June 2024. [Online]. Available: http://arxiv.org/abs/2405.04434

[39] “Mixtral of experts | Mistral AI.” [Online]. Available: https://mistral.ai/news/mixtral-of-experts

[40] “200 languages within a single AI model: A breakthrough in high-quality machine translation.” [Online]. Available: https://ai.meta.com/blog/nllb-200-high-quality-machine-translation/

[41] Lepikhin, H. Lee, Y. Xu, D. Chen, O. Firat, Y. Huang, M. Krikun, N. Shazeer, and Z. Chen, “GShard: Scaling Giant Models with Conditional Computation and Automatic Sharding,” arXiv:2006.16668, June 2020. [Online]. Available: http://arxiv.org/abs/2006.16668

[42] “Coxsackievirus B3 mRNA, complete genome,” Aug. 2018. [Online]. Available: http://www.ncbi.nlm.nih.gov/nuccore/NC_038307.1

[43] A. Green, P. A. Silver, J. J. Collins, and P. Yin, “Toehold Switches: De-Novo-Designed Regulators of Gene Expression,” Cell, vol. 159, no. 4, pp. 925–939, Nov. 2014.

[44] “Synthetic construct monomeric red fluorescent protein gene, complete cds,” Dec. 2004. [Online]. Available: http://www.ncbi.nlm.nih.gov/nuccore/AY678264.1

[45] Jonathan, “jauyong44/vienna-riboswitch,” Jan. 2025. [Online]. Available: https://github.com/jauyong44/vienna-riboswitch

[46] “StandardScaler.” [Online]. Available: https://scikit-learn.org/stable/modules/generated/sklearn.preprocessing.StandardScaler.html

[47] “mixture-of-experts/moe.py at master – davidmrau/mixture-of-experts.” [Online]. Available: https://github.com/davidmrau/mixture-of-experts/blob/master/moe.py

[48] A. Davis and I. Arel, “Low-Rank Approximations for Conditional Feedforward Computation in Deep Neural Networks,” arXiv:1312.4461, Jan. 2014. [Online]. Available: http://arxiv.org/abs/1312.4461

[49] Y. Bengio, N. Léonard, and A. Courville, “Estimating or Propagating Gradients Through Stochastic Neurons for Conditional Computation,” arXiv:1308.3432, Aug. 2013. [Online]. Available: http://arxiv.org/abs/1308.3432

[50] Eigen, M. Ranzato, and I. Sutskever, “Learning Factored Representations in a Deep Mixture of Experts,” arXiv:1312.4314, Mar. 2014. [Online]. Available: http://arxiv.org/abs/1312.4314

[51] L. Denoyer and P. Gallinari, “Deep Sequential Neural Network,” arXiv:1410.0510, Oct. 2014. [Online]. Available: http://arxiv.org/abs/1410.0510

[52] M. I. Jordan and R. A. Jacobs, “Hierarchical Mixtures of Experts and the EM Algorithm,” Neural Computation, vol. 6, no. 2, pp. 181–214, Mar. 1994. doi: 10.1162/neco.1994.6.2.181.

[53] D. E. Rumelhart, G. E. Hinton, and R. J. Williams, “Learning representations by back-propagating errors,” Nature, vol. 323, no. 6088, pp. 533–536, Oct. 1986.

[54] “Linear – PyTorch 2.6 documentation.” [Online]. Available: https://pytorch.org/docs/stable/generated/torch.nn.Linear.html

[55] “ReLU – PyTorch 2.6 documentation.” [Online]. Available: https://pytorch.org/docs/stable/generated/torch.nn.ReLU.html

[56] “torch.nn.functional.softmax – PyTorch 2.6 documentation.” [Online]. Available: https://pytorch.org/docs/stable/generated/torch.nn.functional.softmax.html

[57] A. Lopez-Avila, J. Du, A. Shimary, and Z. Li, “Positional encoding is not the same as context: A study on positional encoding for Sequential recommendation,” arXiv:2405.10436, May 2024. [Online]. Available: http://arxiv.org/abs/2405.10436

[58] V. Passricha and R. K. Aggarwal, “End-to-End Acoustic Modeling Using Convolutional Neural Networks,” in Intelligent Speech Signal Processing, N. Dey, Ed. Academic Press, Jan. 2019, pp. 5–37.

[59] “CrossEntropyLoss – PyTorch 2.6 documentation.” [Online]. Available: https://pytorch.org/docs/stable/generated/torch.nn.CrossEntropyLoss.html

[60] “f1_score.” [Online]. Available: https://scikit-learn.org/stable/modules/generated/sklearn.metrics.f1_score.html

[61] L. Breiman, “Random Forests,” Machine Learning, vol. 45, no. 1, pp. 5–32, Oct. 2001.

[62] J. H. Friedman, “Greedy function approximation: A gradient boosting machine,” The Annals of Statistics, vol. 29, no. 5, pp. 1189–1232, Oct. 2001. doi: 10.1214/aos/1013203451.

[63] L. Khatib, A. Pham, K. Ahmed, and V. S. Frenkel, “Data Centers and Water: Challenges and Solutions for Sustainable Cooling,” J AWWA, vol. 117, pp. 48–53, 2025. doi: 10.1002/awwa.2504.

[64] H. Long, “RiboMoE: An Application of Mixture of Experts on Artificial Riboswitch Classification,” Master of Science thesis, San Jose State University Department of Computer Science, 2025.

[65] L. Breiman, J. H. Friedman, R. A. Olshen, and C. J. Stone, Classification and Regression Trees. Routledge, 2017. doi: 10.1201/9781315139470.

[66] S. Vinga and J. Almeida, “Alignment-free sequence comparison—a review,” Bioinformatics, vol. 19, no. 4, pp. 513–523, 2003.

[67] J. A. Valeri et al., “Sequence-to-function deep learning frameworks for engineered riboregulators,” Nature Communications, vol. 11, art. 5058, 2020. doi: 10.1038/s41467-020-18676-2.

[68] Q. Zhang, S.-R. Jun, M. Leuze, D. Ussery, and I. Nookaew, “Viral Phylogenomics Using an Alignment-Free Method: A Three-Step Approach to Determine Optimal Length of k-mer,” Scientific Reports, vol. 7, art. 40712, Jan. 2017. doi: 10.1038/srep40712.

[69] N. M. Angenent-Mari, A. S. Garruss, L. R. Soenksen, G. Church, and J. J. Collins, “A deep learning approach to programmable RNA switches,” Nature Communications, vol. 11, art. 5057, 2020. doi: 10.1038/s41467-020-18677-1.

[70] “ViennaRNA Web Services,” RNA WebSuite. [Online]. Available: http://rna.tbi.univie.ac.at/ Accessed: Aug. 2023.

[71] L. Breiman, “Random Forests,” Machine Learning, vol. 45, pp. 5–32, 2001. doi: 10.1023/A:1010933404324.

[72] O. Peretz, M. Koren, and O. Koren, “Naive Bayes classifier–An ensemble procedure for recall and precision enrichment,” Engineering Applications of Artificial Intelligence, vol. 136, art. 108972, 2024. doi: 10.1016/j.engappai.2024.108972.

[73] “Escherichia phage MS2 isolate Hu-Sh-YMS-1704-01, complete genome,” Aug. 2021. [Online]. Available: https://www.ncbi.nlm.nih.gov/nuccore/MT830620.1

[74] M. L. Sin, K. E. Mach, P. K. Wong, and J. C. Liao, “Advances and challenges in biosensor-based diagnosis of infectious diseases,” Expert Review of Molecular Diagnostics, vol. 14, no. 2, pp. 225–244, 2014, doi: 10.1586/14737159.2014.888313

[75] Y. Zheng, X. Song, Z. Fredj, S. Bian, and M. Sawan, “Challenges and perspectives of multi-virus biosensing techniques: A review,” Analytica Chimica Acta, vol. 1244, Art. no. 340860, 2023, doi: 10.1016/j.aca.2023.340860

