## Supplementary Information for "Characterization of artificial riboswitches for Coxsackievirus B3 detection using machine learning"

### A Supplementary Information

#### A.1 Feature Statistics

Table A.1: Overall feature statistics computed across the balanced dataset. Values summarize positive and negative constructs together after class balancing.

| Feature | Meaning | Range (min–max) | Mean ± SD |
|---|---|---|---|
| salt_molarity | Monovalent salt concentration used in folding | 0.01–0.01 | $0.01 \pm \sim 0$ (constant) |
| temperature (°C) | Folding temperature | 25.0–25.0 | $25.0 \pm 0$ (constant) |
| trigger_target_binding_percentage | Fraction of target nucleotides paired to the trigger | 0.000–1.000 | $0.621 \pm 0.423$ |
| switch_energy | MFE of the riboswitch (switch) | $-48.17$–$-17.38$ | $-27.75 \pm 3.62$ |
| target_energy | MFE of the target | $-7.66$–0.00 | $-2.70 \pm 2.24$ |
| reporter_energy | MFE of the reporter | $-21{,}474{,}836.48$–$-221.49$ | $-1.074 \times 10^7 \pm 1.074 \times 10^7$ |
| complex_energy | MFE of the whole complex | $-93.14$–$-25.19$ | $-61.19 \pm 19.21$ |
| switch_gc | GC content of the switch | 0.3934–0.5738 | $0.4562 \pm 0.0243$ |
| switch_at | AT content of the switch | 0.4262–0.6066 | $0.5438 \pm 0.0243$ |
| target_gc | GC content of the target | 0.400–0.600 | $0.4724 \pm 0.0397$ |
| target_at | AT content of the target | 0.400–0.600 | $0.5276 \pm 0.0397$ |
| reporter_gc | GC content of the reporter | 0.62025–0.62025 | $0.62025 \pm \sim 0$ (constant) |
| reporter_at | AT content of the reporter | 0.37975–0.37975 | $0.37975 \pm \sim 0$ (constant) |
| complex_gc | GC content of the complex | 0.4012–0.5741 | $0.4617 \pm 0.0211$ |
| complex_at | AT content of the complex | 0.4259–0.5988 | $0.5383 \pm 0.0211$ |
| switch_ensemble_defect | Avg. proportion of mis-paired bases (switch) | 0.0128–0.6486 | $0.1226 \pm 0.0634$ |
| target_ensemble_defect | Ensemble defect (target) | 0.0058–0.3229 | $0.1359 \pm 0.0766$ |
| reporter_ensemble_defect | Ensemble defect (reporter) | 0.0394–0.3468 | $0.1931 \pm 0.1537$ |
| complex_ensemble_defect | Ensemble defect (complex) | 0.0126–0.6513 | $0.0974 \pm 0.0498$ |
| switch_single_strandedness | Avg. unbound likelihood (switch) | 0.6592–0.7901 | $0.7344 \pm 0.0191$ |
| target_single_strandedness | Avg. unbound likelihood (target) | 0.6934–0.9100 | $0.8192 \pm 0.0532$ |
| reporter_single_strandedness | Avg. unbound likelihood (reporter) | 0.6947–1.0000 | $0.8474 \pm 0.1526$ |
| complex_single_strandedness | Avg. unbound likelihood (complex) | 0.6389–0.8002 | $0.7007 \pm 0.0315$ |
| trigger_single_strandedness | Avg. unbound likelihood (trigger) | 0.2330–0.8643 | $0.6420 \pm 0.0850$ |
| rbs_location | Position index of the ribosome binding site | 68.0–75.0 | $71.74 \pm 3.49$ |
| loop_energy | MFE contribution of the loop | 0.0–0.0 | $0.0 \pm 0$ (constant) |

Table A.1 provides the complete descriptive statistics for the structural and thermodynamic features evaluated in the balanced CVB3 dataset. Several features exhibited substantial variation across constructs, whereas others, including `salt_molarity`, temperature, and `loop_energy`, were constant or nearly constant. Class-specific distributions of the most informative features are discussed in Section 3.2 of the main text.

Features computed using ViennaRNA including: trigger-target binding percentage, minimum free energy, GC content, AT content, ensemble defect, single strandness. The values of these features in the positive vs. negative class are shown in the histograms of the Figures A.1 to A.15 below.

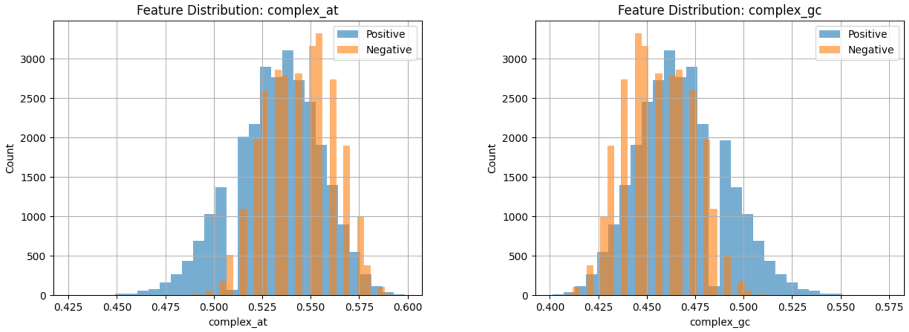


Figure A.1: Left: complex AT content. Right: complex GC content.

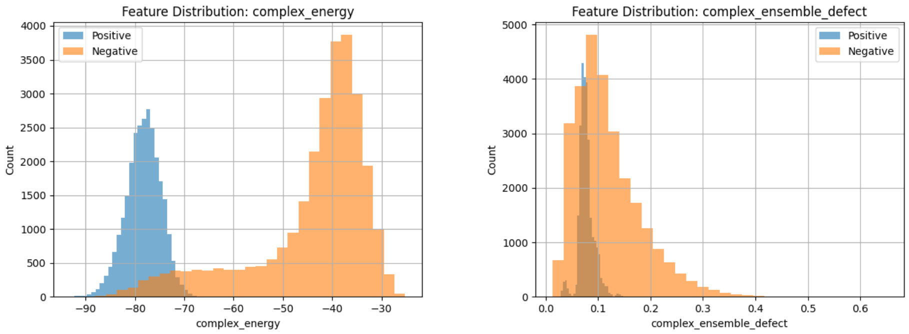


Figure A.2: Left: complex energy. Right: complex ensemble defect.

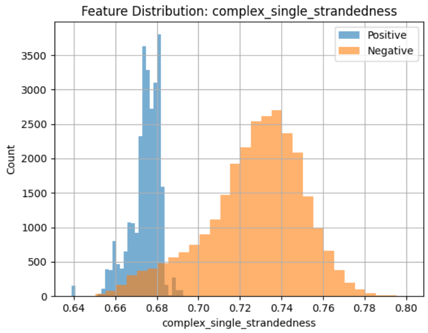


Figure A.3: Complex single strandness

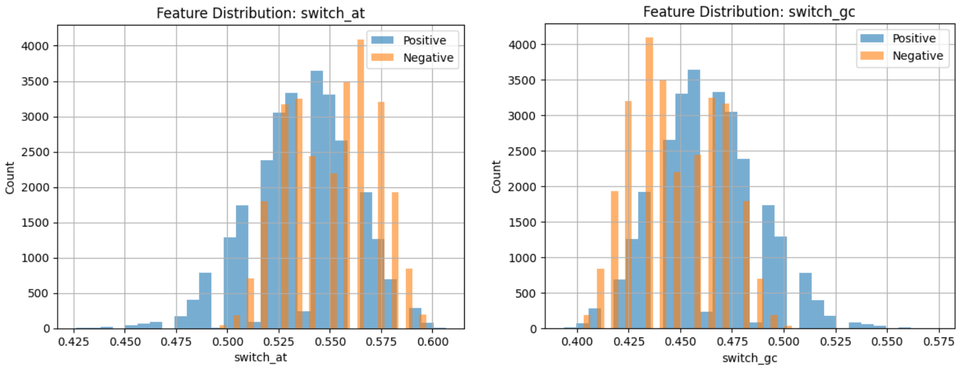


Figure A.4: Left: switch AT content. Right: switch GC content.

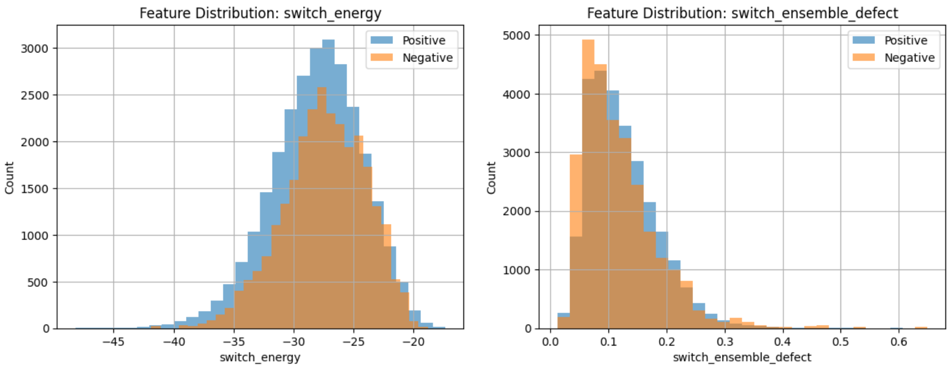


Figure A.5: Left: switch energy. Right: switch ensemble defect.

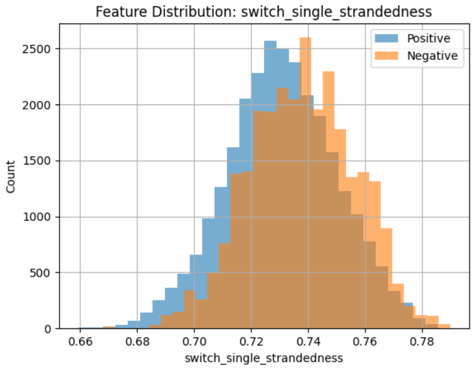


Figure A.6: Switch single strandness

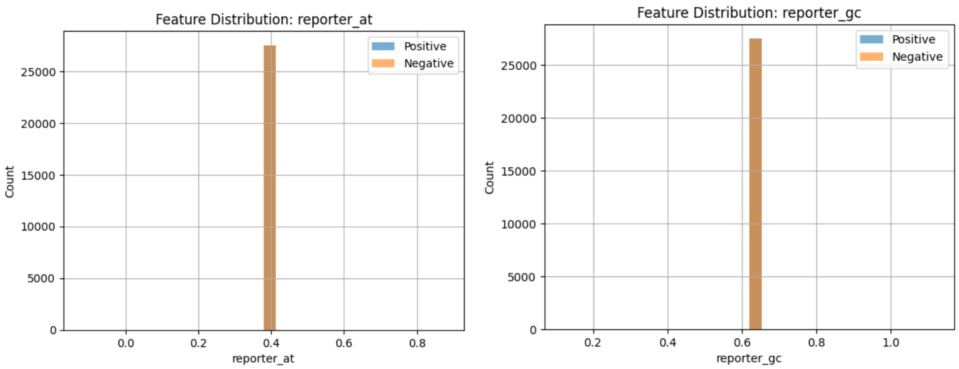


Figure A.7: Left: reporter AT content. Right: reporter GC content.

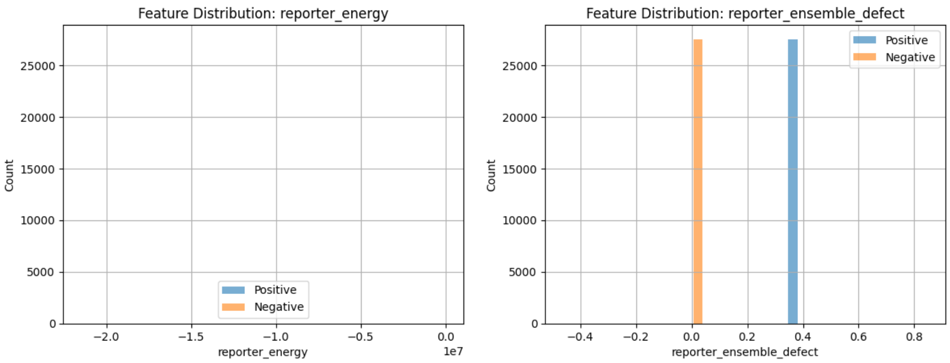


Figure A.8: Left: reporter energy. Right: reporter ensemble.

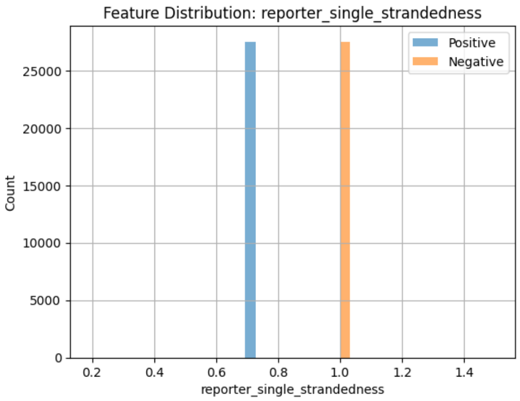


Figure A.9: Reporter single strandness

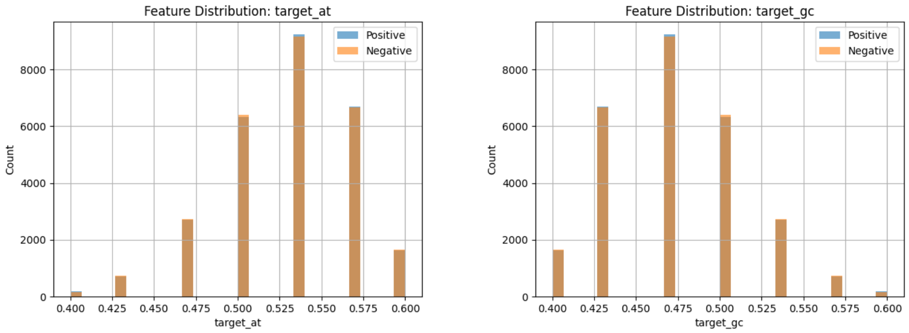


Figure A.10: Left: target AT content. Right: target GC content.

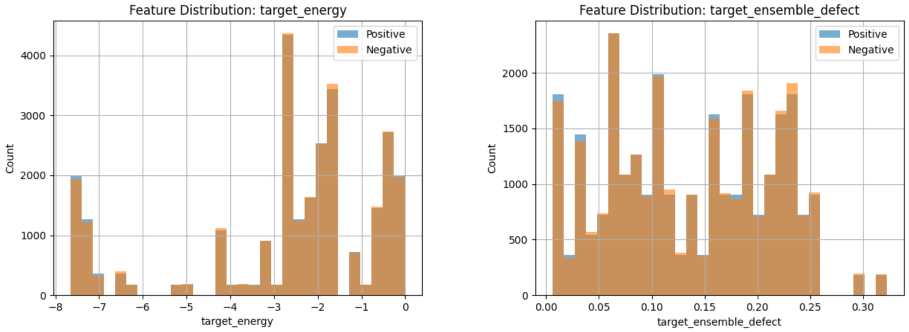


Figure A.11: Left: target energy. Right: target ensemble defect.

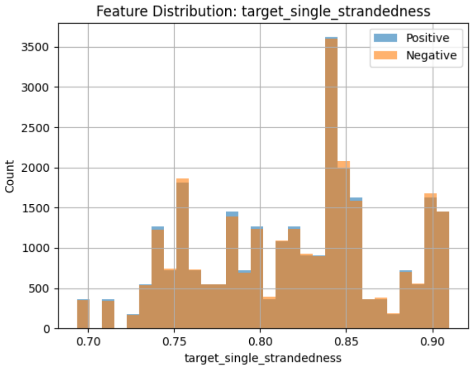


Figure A.12: Target single strandness

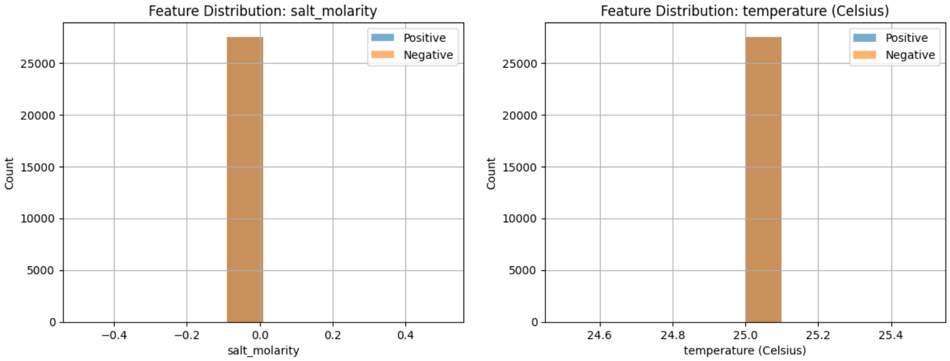


Figure A.13: Left: salt molarity. Right: temperature.

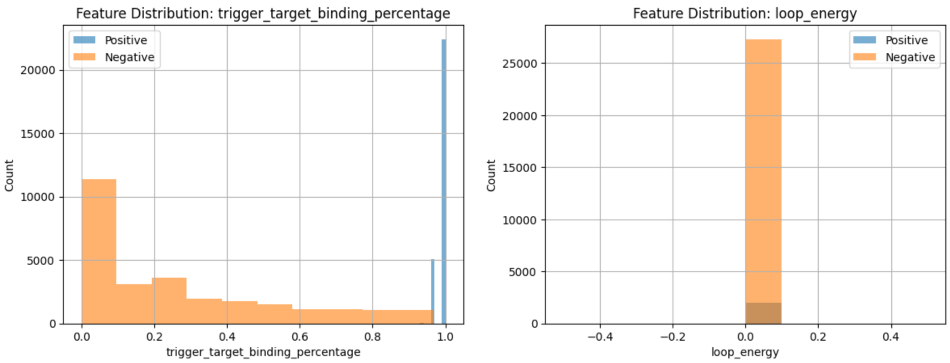


Figure A.14: Left: Trigger target binding percentage. Right: loop energy.

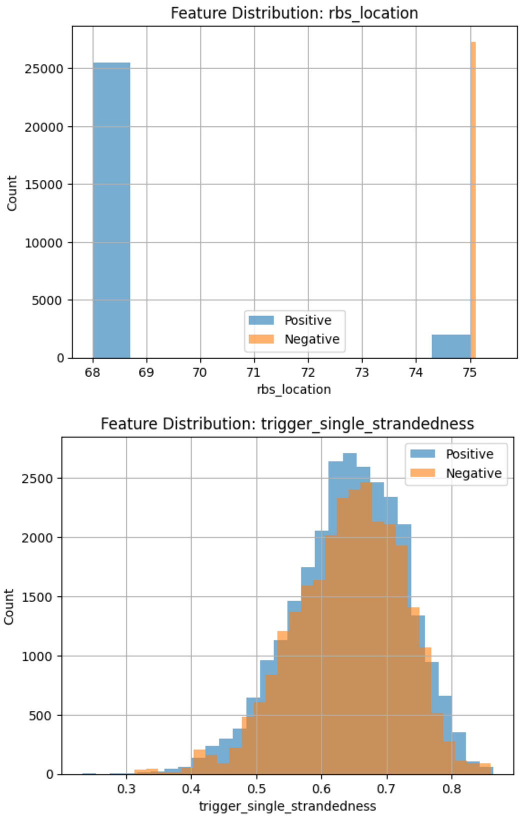


Figure A.15: Top: RBS location. Bottom: Trigger single strandness.

#### A.2 K-mer Analysis

K-mer counts were used as an alignment-free representation of local RNA sequence composition. For each target, trigger, and whole-complex sequence, 3-mer, 4-mer, and 5-mer counts were computed after converting thymine (T) to uracil (U). Sequence-composition patterns were compared across Coxsackievirus B3 (CVB3), MS2, and the balanced positive/negative CVB3 dataset.

For positional analysis, 3-mer frequencies were calculated using a 10-nucleotide sliding window with a 3-nucleotide step size. The five most frequent 3-mers were tracked across window positions. The CVB3 target analysis is presented in Figure 16 of the main text. Additional analyses of the CVB3 trigger and MS2 target and trigger sequences are shown in Figure A.16.

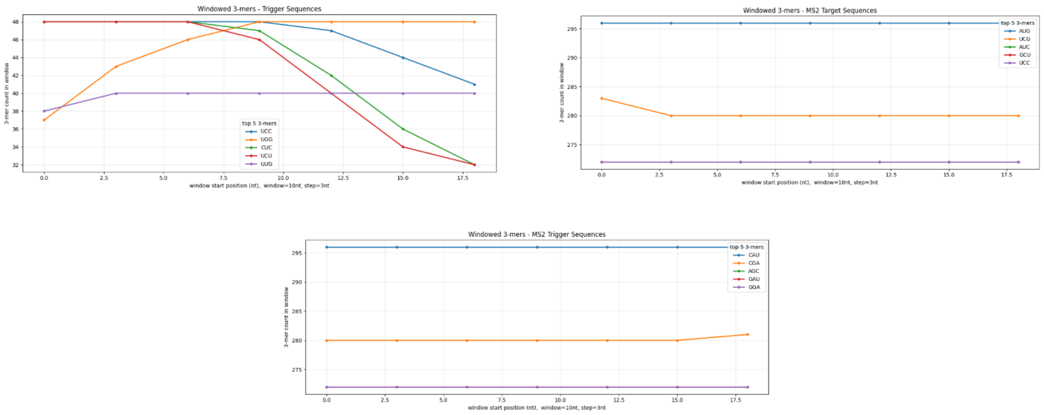


Figure A.16: Windowed 3-mer distributions for the CVB3 trigger and MS2 target and trigger sequences using a 10-nt window and 3-nt step size.

As shown in Figure A.16, the CVB3 trigger sequence exhibited greater positional variation in 3-mer frequencies than the MS2 target and trigger sequences, which showed comparatively stable profiles across window positions. Together with the CVB3 target analysis presented in Figure 16, these results indicate greater positional variation in the CVB3 sequence design space. Subsampling MS2 to match the CVB3 sample size retained a lower flatness score, indicating that the observed difference was not explained by sample size alone. The greater positional variation in CVB3 may reflect sequence-context characteristics of the RNA-dependent RNA polymerase coding region; however, additional codon-position or reading-frame analyses would be required to evaluate this possibility.

Positive-versus-negative sequence differences were evaluated separately using the balanced CVB3 dataset. For each sequence component, differences were calculated as the mean 3-mer count in positive constructs minus the mean count in negative constructs. Positive values therefore indicate enrichment in positive constructs, whereas negative values indicate enrichment in negative constructs.

Table A.2: Largest positive-versus-negative 3-mer differences by sequence component in the balanced CVB3 dataset.

| Sequence Component | Largest Absolute Difference | Dominant 3-mer |
|---|---|---|
| target_sequence | 0.0054 | UUG |
| trigger_sequence | 0.0086 | UCA |
| complex_nucleotides | 1.3604 | UCU |

The target- and trigger-sequence differences were small relative to the whole-complex sequence (Table A.2). The substantially larger difference observed for `complex_nucleotides` indicates that the whole-complex representation contains a stronger class-associated sequence signal than either the target or trigger sequence alone. The corresponding distribution of positive-versus-negative 3-mer differences is shown in Figure 17 of the main text.
